# The 4R Genome Duplication In Salmonine Fishes: Insights From Conserved Non-Coding Elements

**DOI:** 10.1101/2020.09.24.278218

**Authors:** Anibal H. Castillo

## Abstract

Gene and genome duplications are essential processes in evolution. Salmonids are ideal animal model systems to study these processes, as they originated from a tetraploid ancestor. Conserved non-coding elements (CNEs) are of interest because of their highly conserved DNA consensus motifs spanning lineages as diverse and divergent as humans and fish. The main goal of this study is to test CNEs as a tool to study genome duplications and to revisit the “4R” hypothesis and phylogeny of Salmonine fishes (Salmonidae) *Salmo salar*, *Salvelinus alpinus* and *Oncorhynchus mykiss* through the study of copy number and nucleotide variation in six pairs of CNEs. Allele numbers for most CNE sequence pairs are consistent with the 4R hypothesis, as is the symmetric phylogenetic topology shown by some CNE pairs; the estimated date of CNE duplication is consistent with the only reported range of 25-100Mya. However, the phylogenetic relationships within Salmoninae remain unresolved.

## Introduction

Recent advances in genomic approaches are revolutionizing our understanding of chromosomal and genome evolution (Jaillon et al. 2004; Froschauer et al. 2006), gene regulation (Woolfe et al. 2005) and the genetic architecture of phenotypic variation (Mackay 2001). These advances are also broadening our understanding of the patterns of gene and genome duplications (Taylor et al. 2001a,b; Robinson-Rechavi et al. 2001a,b,c; Meyer and Van de Peer 2003; Gregory 2005), for example, by allowing us to infer how many duplication events occurred during a taxon’s evolutionary history and when they occurred (Friedman and Hughes 2001; Froschauer et al. 2006). Genomic analyses in an evolutionary framework can provide insights to answer how many polyploidization events occurred during the evolution of a taxon and understand the timing and reasons behind them (Froschauer et al. 2006).

The idea that gene duplications and polyploidization events are essential processes in vertebrates’ evolution was first proposed in the 1970s by Ohno (1970; see also Allendorf and Thorgaard 1984 and Ohno 1999 for reviews). However, the recent outburst of genomic data generation (Hubbard et al. 2007) due to recent developments in technology has allowed testing established hypotheses in innovative ways.

Among vertebrates, polyploidization is common in fishes, and there are also cases in reptiles and amphibians (Venkatesh 2003; Gregory and Mable 2005). Within fishes, polyploidization has been documented in non-teleosts, such as paddlefish (Polyodontidae), sturgeon (Acipenseridae) and spotted gar (Lepisosteidae) (Allendorf and Thorgaard 1984; Venkatesh 2003), as well as teleosts, such as cyprinids (carps, family Cyprinidae), and the ancestor of the family Salmonidae (Venkatesh 2003; Gregory and Mable 2005). Two whole-genome duplications (WGD) are thought to have occurred in the vertebrate lineage: one before the appearance of the Agnatha, and a second one after the divergence of Agnatha but before the divergence of cartilaginous fishes (Lundin 1993; Furlong and Holland 2002; Moghadam et al. 2005; Van de Peer and Meyer 2005; Froschauer et al. 2006; Sémon and Wolfe 2007). This idea, known as the “2R Hypothesis” (Larhammar et al. 2002; see Kasahara 2007 for a review), is now widely accepted, although until very recently, it was still controversial (Robinson-Rechavi et al. 2001a,b; Taylor et al. 2001c; Jaillon et al. 2004; Stellwag 2004). An additional genome duplication is thought to have occurred in the ancestral fish lineage before the emergence of teleosts (“3R Hypothesis”), approximately 320-450 million years ago (Mya) (Stellwag 2004; Van de Peer and Meyer 2005; Froschauer et al. 2006). Moreover, an independent tetraploidization event is thought to have occurred in the ancestor of Salmonids approximately 25-100 Mya (the “4R Hypothesis”) (Allendorf and Thorgaard 1984). After these WGDs, tandem or segmental duplications of individual genes or groups of genes might have taken place at different rates in different vertebrate lineages. Following duplication, possible scenarios involve pseudogenization of one gene copy by the accumulation of deleterious mutations, the evolution of one copy to encode a protein with a new biological function and retention of both derived genes to perform the function of the ancestral gene (i.e. sub-functionalization, Lynch and Force 2000a,b; Robinson-Rechavi et al. 2001c; Taylor and Raes 2005; Froschauer et al. 2006; Zhu et al. 2007, see Zhang 2003 for a review).

There has been a long debate about the relative role of the different processes explaining the differences in gene copy number. For example, Robinson-Rechavi et al. (2001a) suggest that the increase in gene copies in teleosts might be due to gene families’ expansions rather than an ancient WGD. However, many duplicated genes in distantly related fish species such as pufferfish and zebrafish map to the same paralogous chromosomes, suggesting a WGD event has occurred (Venkatesh 2003). In turn, the evolution of some genes can be explained by both processes (Kawasaki et al. 2007). Conversely, some duplicated genes in zebrafish do not show a phylogenetic topology consistent with the WGD hypothesis. Many processes can explain an alternative topology, such as selective sweeps, absence of sequence variation, different mutation rates between duplicates, acquisition of a new function or subfunctionalization, among others (Lynch and Force 2000a,b; Friedman and Hughes 2001; Taylor and Raes 2005). There is also the possibility that individual gene duplications may have played a significant role in shaping these species’ genetic architecture (Massingham et al. 2001; Robinson-Rechavi et al. 2001a,d; Kellogg 2003a,b).

There are four ways in which genomic DNA can duplicate. These are tandem repeats, transposition duplications, chromosomal segment duplications, and entire genome duplications (Li et al. 2003). As complex phenomena, WGD are usually studied through many approaches. It has been argued that the most rigorous way of testing genome doublings is identifying paralogous chromosomal regions or block duplications (Lundin et al. 2003). For example, if a descendant species has twice the number of chromosome arms than an ancestral grouping and are identified as paralogous, this would infer a WGD. Once a statistically significant number of pairs of homologous regions is identified, a WGD can be inferred (Van de Peer and Meyer 2005). This approach requires a map-based dataset, including markers’ absolute or relative position in the species’ genome sequence. In addition to these processes, there are also “hidden duplications”, those very degenerated (i.e. where substantial gene loss has occurred) that can only be diagnosed through comparison with a third genome (Van de Peer and Meyer 2005).

Although capable of providing excellent insight into the study of genome duplications, the approaches mentioned above are not devoid of problems. They rely on extensive datasets (e.g. many pairs of markers) that are inherently expensive and require mapping information contingent upon the availability of detectable polymorphisms (i.e. variable markers). Moreover, for some groups of undeniable biological interest, like the viscacha rat *Tympanoctomys barrerae*, the first mammal species ever reported as a polyploid (Gallardo 1999; 2003; 2004; 2006), it is unlikely that the required economic effort will be made in the foreseeable future, in part because of the species’ lack of commercial interest.

The 4R genome duplication is well established in Salmonids (Allendorf and Thorgaard 1984; Phillips and Ráb 2001; Moghadam et al. 2005; Gregory and Mable 2005), although not extensively characterized. The genetic architecture of the genomes in species within the subfamily Salmoninae is partially understood, as genetic maps are under construction (Danzmann et al. 2005) and genome sequencing projects for rainbow trout and Atlantic salmon are underway (genamics.com). Therefore, Salmonines offer an excellent case to test the predictions of a WGD by investigating if the effects of this duplication can be recovered on specific markers. If so, this method can be used in the groups mentioned before, where genome studies are of great biological interest, but more arduous efforts are unlikely to shed light on genome duplications.

One exciting set of sequences is the highly conserved non-coding sequences (Conserved Non-coding Elements, hereafter “CNEs”). In a comparative analysis of the genome sequences of humans *(Homo sapiens)* and the freshwater green-spotted pufferfish (*Tetraodon nigroviridis*), Woolfe et al. (2005) identified 1373 CNEs. Most of these sequences appeared to regulate developmental genes (Woolfe et al. 2005). Moreover, these CNEs appear to be unique to vertebrates, as none of them has so far been found in invertebrates.

The discovery of CNEs provides an opportunity to characterize CNEs as a tool to investigate WGDs and revisit the 4R whole-genome hypothesis in Salmonine fishes. In particular, it will be possible to test whether the number of copies present in the genome of Salmonids arose from individual sequence duplications or corresponds to a WGD and whether the topology of a phylogenetic tree of these sequences is consistent with the Salmonines’ known history of tetraploidization. If this latter hypothesis holds, loci that have multiple copies in Salmonine genomes should have a symmetric gene tree (Friedman and Hughes 2001), since alleles from each putative duplicated CNE are expected to be more related to each other than they are to alleles from their homeologue CNE duplicate (Figure 1*a*). If the pattern of homology of the CNEs is not due to WGDs but, for instance, due to segmental duplications, then the branching pattern will reflect the individual duplication events (Figure 1*b*). Moreover, for those loci that show a topology consistent with a WGD, an attempt can be made to date such duplication, and the estimated date can be compared and contrasted with the one reported in the literature, 25-100Mya (Allendorf and Thorgaard 1984). Given that this wide range is the only one available for the genome duplication in the salmonid fishes’ ancestor, this issue awaits more precise resolution (Gregory and Mable 2005).

**Figure 1.**
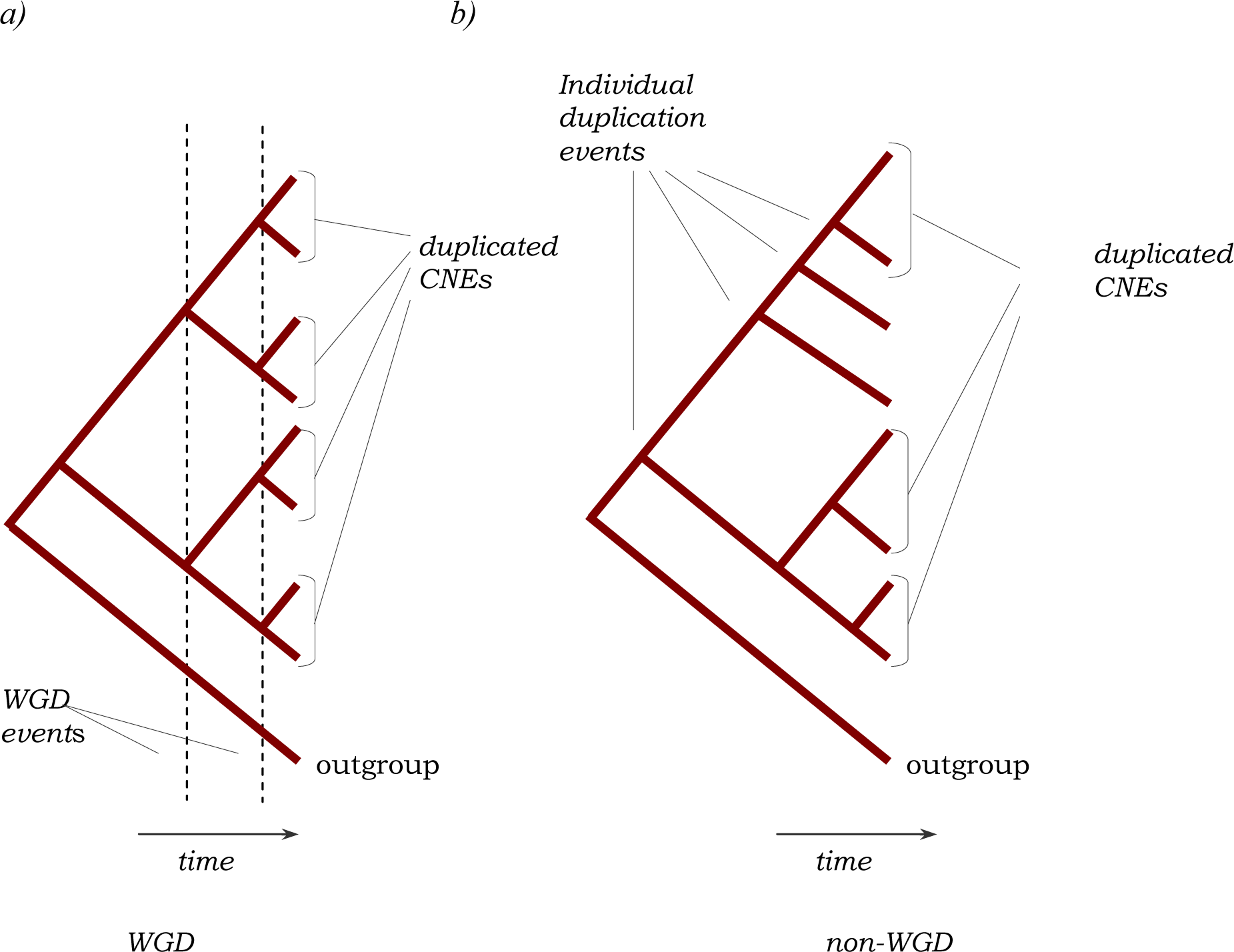
Possible phylogenetic topologies for the CNEs. a) Expected phylogenetic pattern due to a Whole Genome Duplication (WGD). The duplicated CNEs cluster in pairs and their common ancestors date back to the same instant in the past, the time of the WGD event. b) If the pattern of homology of the CNEs is not due to WGDs, but for example, to segmental duplication, the phylogenetic pattern will reflect the individual duplication events.

When phylogenies are used to date duplication events such as the 2R, 3R or 4R hypothesis, the sequences to be included in the analyses should have similar mutation rates, which means the molecular clock hypothesis holds (Taylor et al. 2003). This hypothesis can be tested with a Maximum Likelihood approach through a likelihood ratio test (LRT, Felsenstein 1981). If this test fails to reject the molecular clock hypothesis, it would suggest rate homogeneity across the sequences (Liò and Goldman 1998); therefore, nucleotide divergence can be considered proportional to time. In contrast, if the molecular clock hypothesis does not hold, a relaxed molecular clock can be used, that is, one that accounts for lineage-specific rate heterogeneity (Drummond et al. 2006). Furthermore, through the use of calibration points such as fossils, an age can be estimated for the duplication.

Gene trees of CNEs can also determine phylogenetic relationships of groups whose affinities have been controversial, such as Salmonids. The phylogenetic relationships within the subfamily Salmoninae have been extensively studied (Fulton and Crespi 2003 and references therein). In this regard, the monophyly of particular groups has been established, like the trout genus *Oncorhynchus* (Fulton and Crespi 2003). However, controversy still exists as to whether the genus *Salmo* is more closely related to the genus *Oncorhynchus* (Phillips and Pleyte 1991; Stearley and Smith 1993; Murata et al. 1996; Phillips and Oakley 1997) or *Salvelinus* (Oakley and Phillips 1999; Crespi and Fulton 2003; Kinnison and Hendry 2003). On a related note, addressing these questions with CNE sequences will help determine if they are an appropriate phylogenetic tool.

Any conclusion arising from model-based phylogenetic analyses of salmonid taxa must be based on statistically supported models. Using arbitrarily chosen models might yield results that lend support to different hypotheses (Posada and Crandall 2001a,b). This issue can be addressed, among others, by a test like the LRT (Felsenstein 1981; Posada and Crandall 2001a,b; Bos and Posada 2005) that evaluates the compromise between increasing the complexity of a model and maximizing the accuracy of parameter estimation (Posada and Crandall 1998). LRTs are based on comparisons of estimations of a given tree topology’s likelihood according to different molecular evolution models. Therefore, they can be statistically compared, considering the simpler model as a null hypothesis (H0) and the more complex one as the alternative hypothesis (H1).

The number of copies and sequence variation was studied in six pairs of CNEs in three species of salmonid fishes to test four hypotheses:

1. The allele number for a given CNE family will be consistent with the 4R hypothesis. The 4R hypothesis predicts that Salmonids will show twice as many copies as in other teleost fishes such as zebrafish. If three or more core sequences were found among many clones from one fish, it would be strong evidence that there is more than one gene region for this sequence in the genome. Particularly, those loci present as a single copy in zebrafish are expected to show up to four alleles maximum in Salmonines, assuming that each duplicate locus generates two distinct alleles (Figure 2). Similarly, for locus pairs present as duplicated pairs within zebrafish, it might be possible to detect eight distinct allelic copies in the 4R derivative Salmonines.
2. Members of a CNE family will show a symmetric phylogenetic topology consistent with the 4R hypothesis in Salmonine fishes. Those loci that have been duplicated in Salmonines are expected to show symmetric trees, with two basal clades that will include alleles from each of the three species of Salmoninae.
3. The date of inferred CNE duplications will be consistent with the range of 25-100Mya (Allendorf and Thorgaard1984). It is also possible that CNE data can refine this estimate by narrowing the plausible range of times.
4. Assessing the phylogenetic relationships of the three genera of the subfamily Salmoninae with independent loci will confirm the findings by Crespi and Fulton (2003) that the genera *Oncorhynchus* and *Salvelinus* are sister groups.

**Figure 2.**
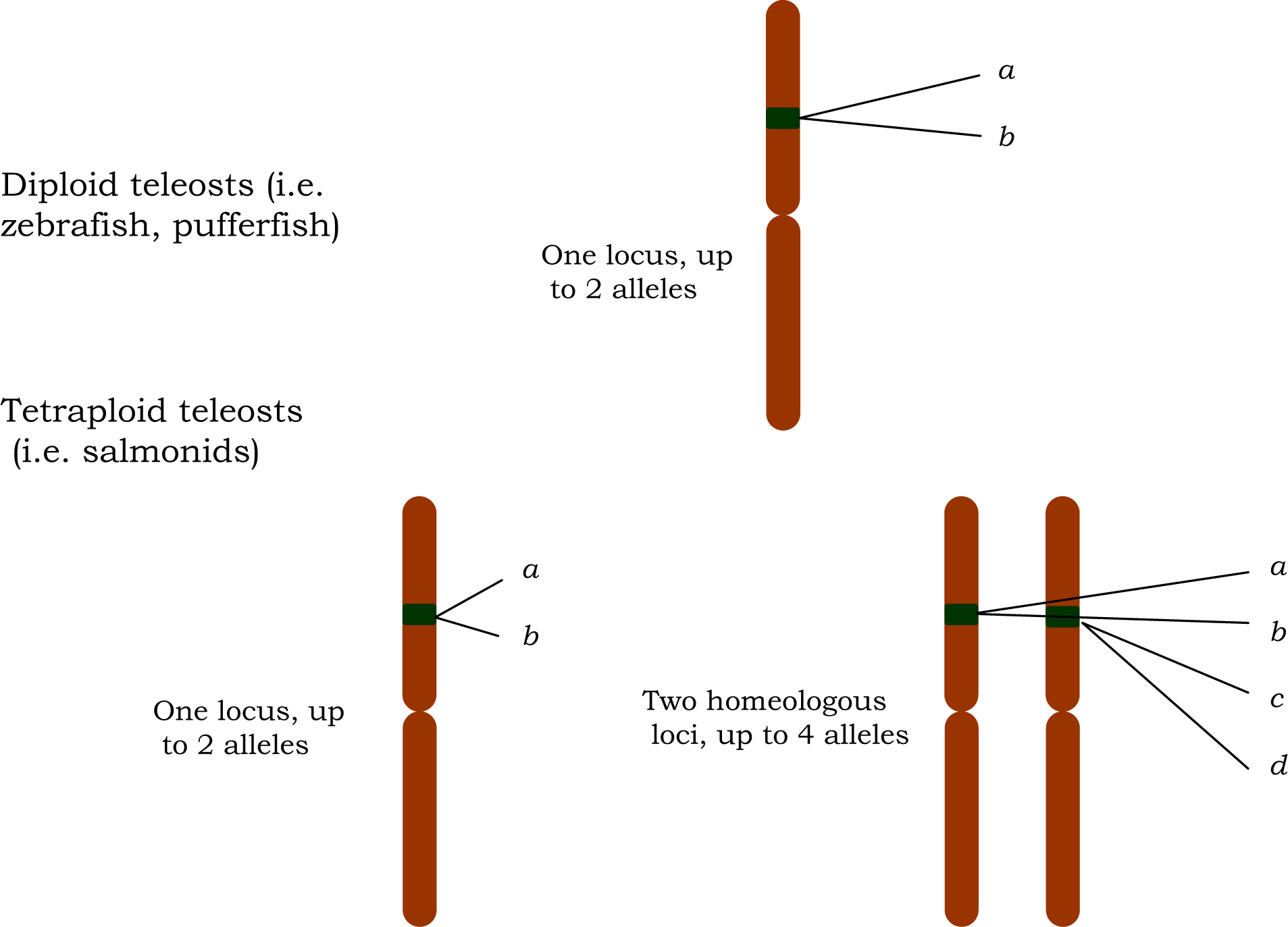
Number of expected alleles in a locus that is single copy in diploid teleosts. If the locus is also single copy in 4R derivative teleost polyploids (i.e. Salmonines), they are expected to show up to two alleles. If it is duplicated, up to four alleles are expected.

Given that several CNEs map close to each other in the human genome and are relatively conserved (Woolfe et al. 2005), we amplified a DNA segment containing two syntenic CNEs in Salmonines (Figure 3). To find nucleotide variation useful for phylogenetic analyses, we chose CNEs physically close to each other, so the intervening region between them and a partial section of each flanking CNE could be amplified in a single polymerase chain reaction (PCR) product, subsequently cloned and sequenced in a single sequencing reaction.

**Figure 3.**
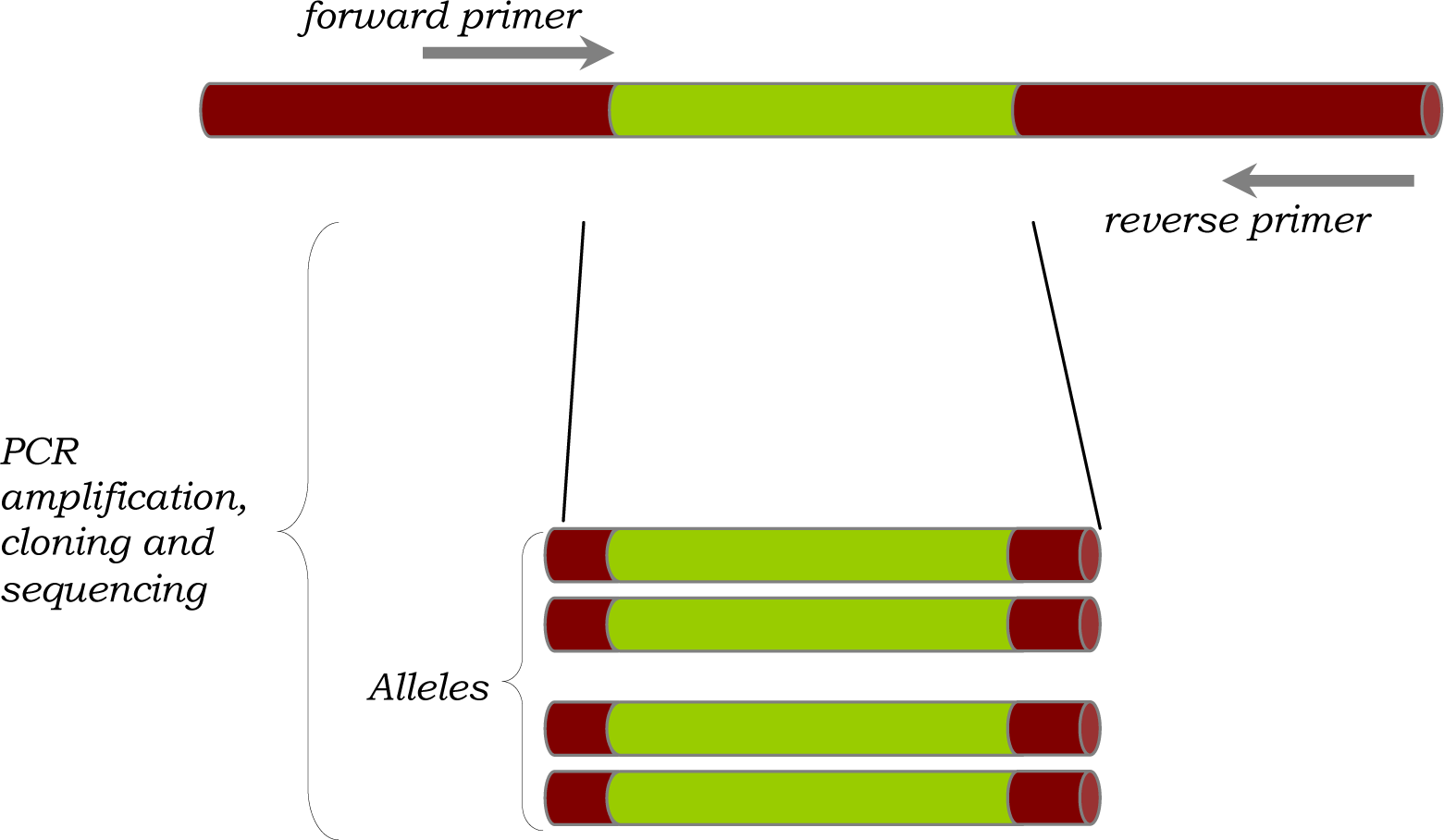
Experimental design for the PCR amplification, cloning and sequencing of a DNA segment containing two CNEs. If three or more core sequences are found among a group of clones from one single fish, it is strong evidence that there is more than one gene region for this sequence in the genome. Given that Salmonines are 4R derivative polyploids, it may be expected to find up to four unique sequences, or alleles, derived from each single copy gene in a 3R species, and up to eight alleles for each duplicated locus pair in a 3R species.

## Materials and Methods

### Loci targeted

The following six pairs of CNEs were amplified: CNE7060-7061, CNE7061-7063, CNE6699-6700, CNE6700-6697, CNE6820-6816 and CNE6739-6741. These loci correspond to partial sequences plus intervening regions of the sequences under GenBank accession numbers CR847060-CR847061, CR847061-CR847063, CR846699-CR846700, CR846700-CR846697, CR846820-CR846816 and CR846739-CR846741 respectively, as annotated in Ensembl database (available at www.ensembl.org/index.html; Hubbard et al. 2007) for teleost fishes zebrafish (*Danio rerio*), green-spotted freshwater pufferfish (*Tetraodon nigroviridis*) and Japanese pufferfish (*Fugu rubripes*). See Table 1 for a description of these loci’s location in the genomes of *Tetraodon* and zebrafish.

**Table 1.** Location of the CNEs flanking loci CNE7060-7061, CNE7061-7063, CNE6699- 6700, CNE6700-6697, CNE6820-6816 and CNE6739-6741 in the genomes of *Tetraodon nigroviridis* (version 47.1i) and *Danio rerio* (Zv7). Data taken from the Ensembl database (release 47, Hubbard et al. 2007).

| Accession number | LG in<br>*Tetraodon* | LG in<br>zebrafish | Duplicates in zebrafish |  |
| --- | --- | --- | --- | --- |
|  |  |  | Start | End |
| CR847060 | Tn17 | Chr:13 | 23149301 | 23149466 |
| CR847061 | Tn17 | Chr:13 | 23148546 | 23148688 |
| CR847063 | Tn17 | Chr:13 | 23148334 | 23148432 |
| CR846699 | Tn3 | Chr:1 | 50568525 | 50568551 |
|  |  | Chr:3 | 41259925 | 41260084 |
| CR846700 | Tn3 | Chr:1 | 3808874 | 3808933 |
|  |  | Chr:3 | 41260308 | 41260422 |
| CR846697 | Tn3 | Chr:1 | 3808978 | 3809248 |
|  |  | Chr:3 | 41260467 | 41260797 |
| CR846820 | Tn1 | Chr:22 | 16338638 | 16338811 |
| CR846816 | Tn1 | Chr:22 | 16338421 | 16338526 |
| CR846739 | Tn8 | Chr:16 | 2332398 | 2332699 |
| CR846741 | Tn8 | Chr:16 | 2332805 | 2332913 |
|  |  | Chr:19 | 12795551 | 12795610 |

The sequences used in this study were chosen according to the putative number of copies that exist in the zebrafish and pufferfish genomes (Table 1):

1. CNE7060-7061, CNE7061-7063 and CNE6820-6816 have been retained as a single copy in both pufferfish and zebrafish.
2. Two loci, CNE6699-6700 and CNE6700-6697, composed of CNEs CR846697, CR846700 and CR846699, are retained as duplicates within zebrafish and map to different linkage groups (LGs).
3. Finally, in CNE6739-6741, its first CNE, CR846739, has been retained as a single copy in the zebrafish’ genome, while the second CNE, CR846741, maps to two different LGs.

In both pufferfish and zebrafish, CNE7060-7061 and CNE7061-7063 and CNE6699-6700 and CNE6700-6697 are adjacent in the same LG, hundreds of base pairs apart (Table 1).

### Species examined

Species of the three genera that comprise the subfamily Salmoninae, rainbow trout *(Oncorhynchus mykiss),* Atlantic salmon *(Salmo salar)* and arctic charr *(Salvelinus alpinus)* were used in this study. The samples were from mapping panels used to construct genetic linkage maps (Jackson et al. 1998; Woram et al. 2003). Nucleotide sequences from zebrafish, green-spotted freshwater pufferfish and Japanese pufferfish were downloaded from the Ensembl database (Hubbard et al. 2007) and used as outgroup taxa in the phylogenetic analyses.

### Molecular analysis

Primers (Table 2) were designed based on nucleotide sequence alignments of zebrafish, the green-spotted pufferfish and the Japanese pufferfish from the Ensembl database, with the assistance of Primer3 (v. 0.4.0, available at fokker.wi.mit.edu/cgi-bin/primer3/primer3_www.cgi). Primers suggested by this software were sometimes modified to ensure that the primer’s last nucleotide was a cytosine or guanosine, thus providing stronger annealing to the template DNA, therefore enhancing PCR performance. Primer names were designed by removing the first four characters from the corresponding CNEs’ GenBank accession number (e.g., “CR84”) and adding “CNE” instead.

**Table 2.**
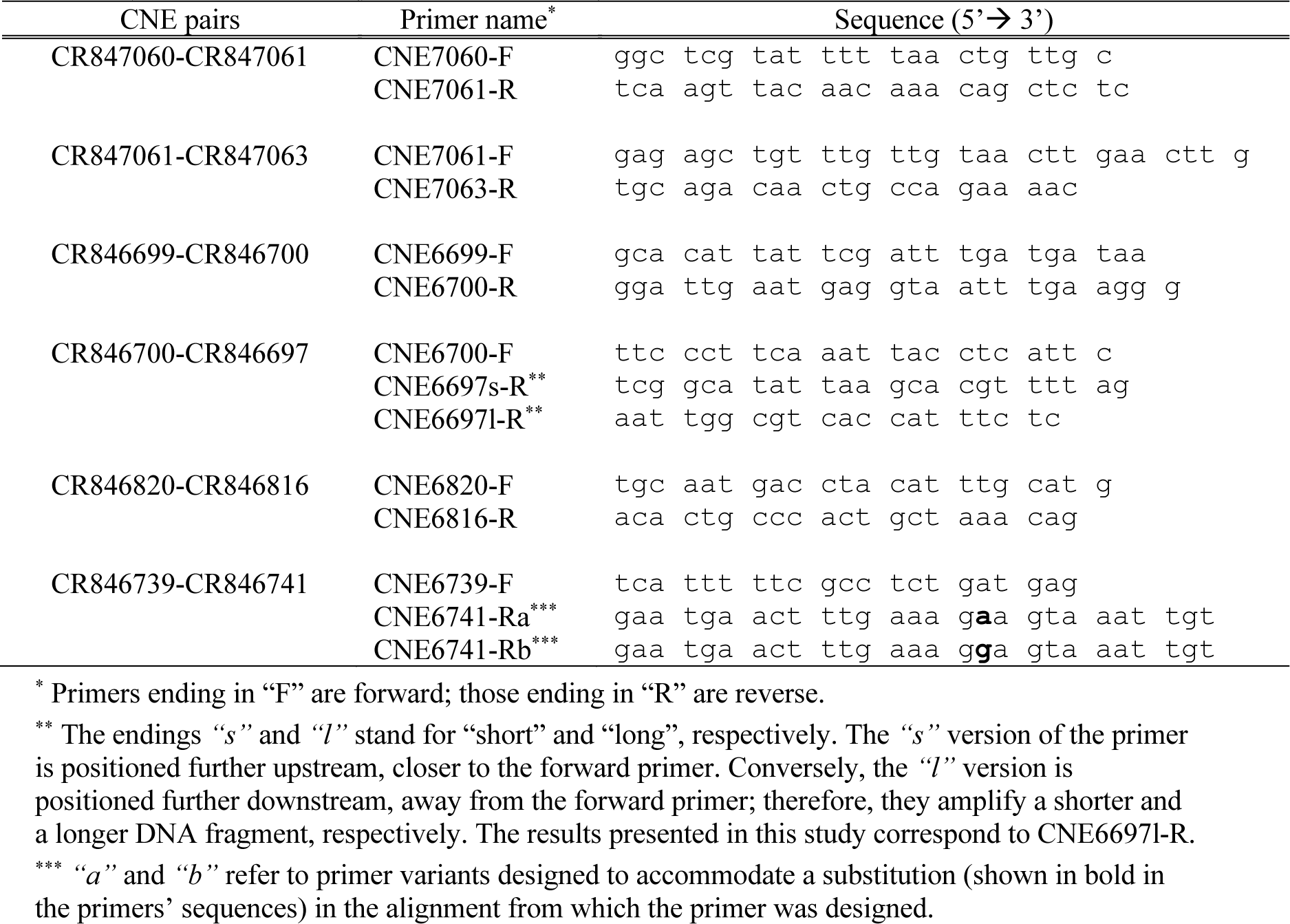
Primers used to amplify the six CNE pairs included in this study. Their design was based on nucleotide sequence alignments of *Danio rerio*, *Tetraodon nigroviridis,* and *Fugu rubripes* from the Ensembl database (Hubbard et al. 2007).

PCR amplifications were carried out in reaction volumes of 11μl-50μl containing 0.02 U/μl of *Taq* polymerase (Life Technologies, Carlsbad, California), 3pg/μl of DNA solution from one single individual, 2.0 mM MgCl2, 0.50μM of each primer and 0.8mM of dNTPs (0.2mM each). Amplification conditions were the following: polymerase activation at 95°C for 5 min, 30 cycles alternating denaturation at 95°C for 30sec, annealing at 48-58°C for 30sec and extension at 72°C for 1 min and 30sec followed by a final extension at 72°C for 5 min.

Aliquots (2μl) of PCR products were run for 20 minutes at 80 volts in 2% agarose gels stained with GelRed^TM^ (Biotium) and visualized in a UV light transilluminator Alpha Imager 3400 Documentation and Analysis System (Alpha Innotech, San Leandro, CA). Each remaining product was purified using Wizard® PCR Preps DNA Purification System (Promega). PCR products were subsequently cloned into pGEM®-T Easy Vector plasmids in a JM109 High-Efficiency Competent Cells (Qiagen) strain of *Escherichia coli*. These plasmids were purified with Plasmid DNA Purification Minipreps (Qiagen), used as a template in cycle sequencing reactions and run on 4% denaturing polyacrylamide gels in an ABI 377 automated sequencer (Applied Biosystems, Foster City, California) using a Big-Dye Terminator Cycle Sequencing kit (Applied Biosystems, USA). Ten to 13 clones were sequenced per locus per species. Only sequences that were different from each other were kept; for those alleles sequenced more than once, extra copies were removed from subsequent analyses. Sequences were aligned using DIALIGN 2 (Morgenstern 1999) and ClustalX (Version 1.81—Thompson et al. 1997, see Appendix I).

### Phylogenetic analyses

Three approaches were employed in the phylogenetic analyses: Bayesian Analysis (BA), Maximum Likelihood (ML) and Maximum Parsimony (MP). BA was performed using MrBayes (v 3.1.2., Huelsenbeck and Ronquist 2001). For this purpose, the Bayesian Information Criterion (BIC, Schwarz 1978) was used to choose a molecular evolution model. This method aims at resolving the compromise between the complexity of the models and the accuracy of parameter estimation. In this regard, 56 models of molecular evolution were assessed with Modeltest (v3.7, Posada and Crandall 1998), and their parameters were estimated using PAUP* (v4.0b10, Swofford 2003). These parameter estimates were used as priors in the Bayesian analyses. The nodes’ posterior probabilities were used to measure their statistical support and were estimated by running four Monte Carlo Markov Chains for one to two million generations to ensure convergence of the parameter estimates. In this regard, only the last 75% of the generations were retained.

Moreover, the same set of 56 models of molecular evolution was also assessed according to two additional criteria: Hierarchical Likelihood Ratio Tests (hLRTs, Felsenstein 1981) and Akaike Information Criterion (AIC, Akaike 1974). Like BIC, these methods aim to resolve the trade-off between the model complexity and parameter estimates’ accuracy (see Posada and Buckley 2004). This hierarchy of nested models can be seen as different hypotheses being tested one at a time. The ability of LRTs to falsify a null hypothesis might depend on the order in which these hypotheses are tested (Posada and Crandall 2001a). Therefore, four different hierarchies (sequences of tests) were used to test the different hypotheses that comprise molecular evolution models, as implemented in MrModelTest (v2.2, Nylander 2004).

The selected models were used to perform a heuristic search (1000 replicates) according to the ML criterion. Furthermore, the most parsimonious trees were obtained using PAUP* (v4.0b10, Swofford 2003), with equal weights for all sites, a heuristic search (1000 replicates) steepest descent option, random additions of taxa and Tree Bisection-Reconnection swapping algorithm. Consensus trees (50% majority rule) of the most parsimonious trees were obtained.

The molecular clock hypothesis was tested for the six datasets. Once a molecular evolution model was chosen, the molecular clock was enforced. The likelihood of the model was recalculated, considered the null hypothesis, and tested against the same model without enforcing a molecular clock (Posada and Crandall 1998) using LRT. The number of degrees of freedom equals the number of taxa minus two (Felsenstein 1981).

In order to date the nodes potentially corresponding to genome duplications, calibration methods were used. In cases where the molecular clock hypothesis was not rejected, a strict molecular clock was enforced. In contrast, in cases where the molecular clock hypothesis was rejected, a relaxed molecular clock was enforced. Nodes including only rainbow trout clones were considered representatives of the genus *Oncorhynchus’ origin* and were assigned an age of six million years (My) (Nelson 2006, p. 202). Clades comprising the genera *Salmo*, *Salvelinus* and *Oncorhynchus* were considered representatives of the origin of the subfamily Salmoninae and were assigned an age of 20My (Groot 1996). All fossil ages were assigned an arbitrary error of ±0.5My. The analyses were performed using BEAST (v1.4, Drummond and Rambaut 2007), and the results were visualized using Tracer (v1.3, Rambaut and Drummond 2007).

## Results

### Intraspecific variation and models of molecular evolution

For loci that have been retained as a single copy in zebrafish, namely CNE7060-7061, CNE7061-7063 and CNE6820-6816, the number of different clones obtained in Salmonines for each locus ranged from six to ten. In particular, CNE7060-7061 resulted in eight clones for rainbow trout and arctic charr and nine clones for Atlantic salmon, whereas CNE7061-7063 showed nine clones for rainbow trout, six clones for Atlantic salmon and ten clones for arctic charr. In locus CNE6820-6816, five different clones were detected in each species. In contrast, loci CNE6699-6700, CNE6700-6699 and CNE6739-6741, which have been retained as double copies in zebrafish, yielded between eight to 11 different clones in Salmonines. In CNE6699- 6700, eight different clones were obtained for each species, and CNE6700-6699 resulted in 11 clones for rainbow trout and Atlantic salmon, and nine clones for arctic charr. For CNE6739- 6741, the results were nine clones for rainbow trout and Atlantic salmon, and seven clones for arctic charr.

The results of model tests are summarized in Table 4. Alternative methods to choose models of molecular evolution, AIC and hLRTs yielded similar results. Four models explain the patterns of molecular evolution of the different loci used in this study. The substitution pattern in CNE6739-6741 can be explained by the model K80 (Kimura 1980) plus a gamma distribution of variable sites. This model includes equal nucleotide frequencies, one parameter to describe mutations both between purines and between pyrimidines (transitions), and another parameter for mutations between nucleotides of different types, purine-pyrimidine (transversions). The molecular evolutionary features of CNE6820-6816 and CNE6699-6700, on the other hand, are best explained by the model HKY85 (Hasegawa et al. 1985). This model differs from K80 in that it incorporates different nucleotide frequencies; therefore, there is a deviation from the null assumption that nucleotide frequencies are 0.25 each.

**Table 3.**
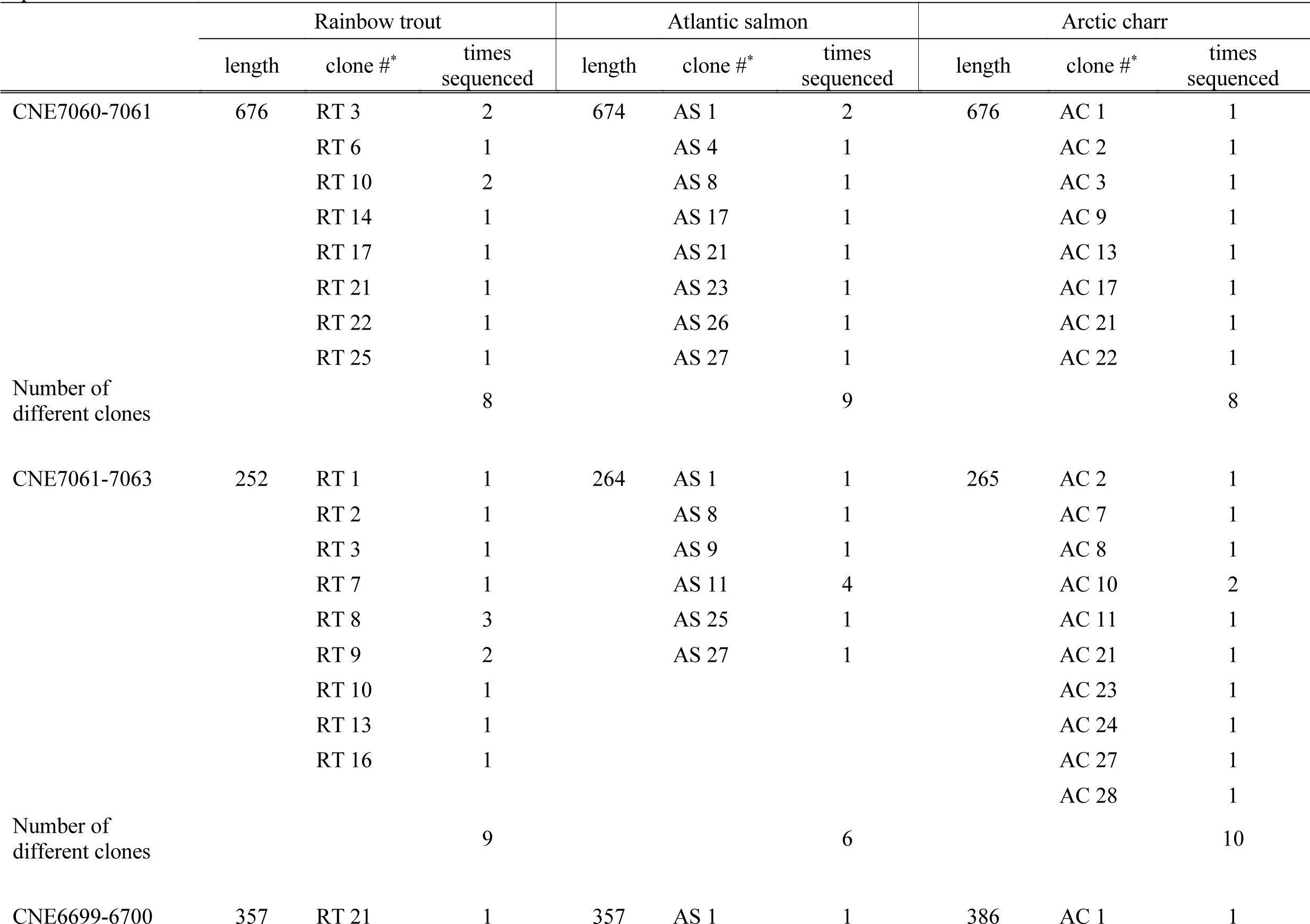
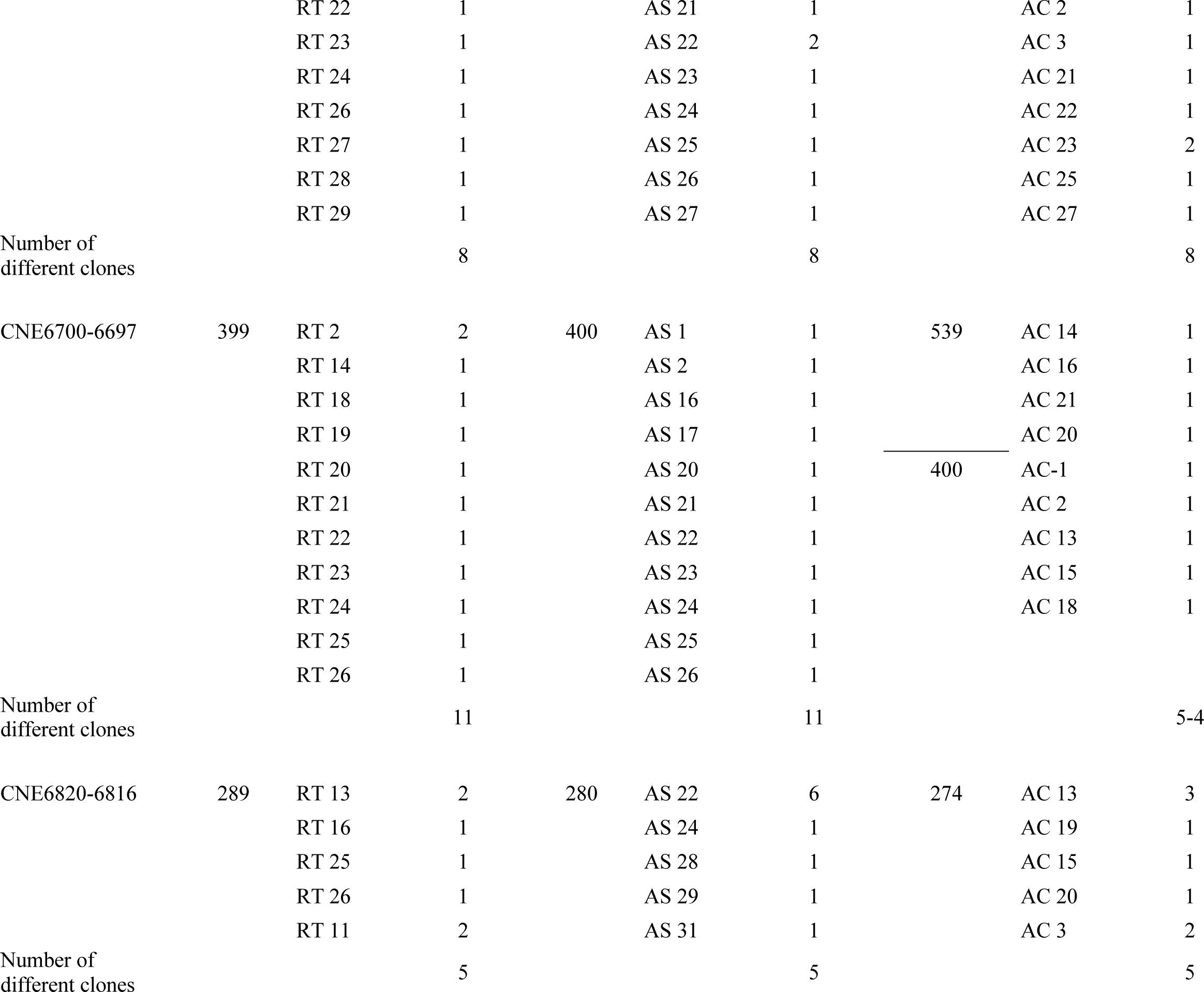
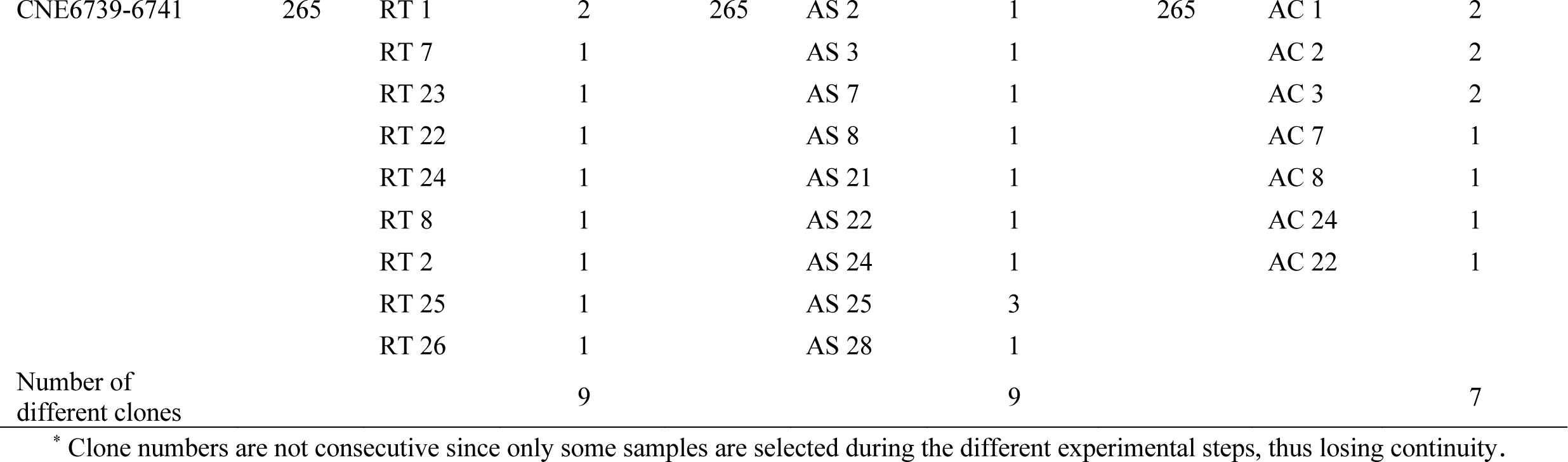
Length (bp) of the CNE pairs plus intervening regions in the three species of Salmonines, rainbow trout (*Oncorhynchus mykiss*), Atlantic salmon (*Salmo salar*) and Arctic charr (*Salvelinus alpinus*) included in this study. The numbers of different clones obtained for each species are indicated.

**Table 4.** Models of molecular evolution selected for each pair of CNEs. TVM = Transversional model; HKY85 = Hasegawa, Kishino and Yano (1985); TrN = Tamura and Nei (1993); K80 Kimura (1980); G = gamma distribution of variable sites; I = proportion of invariable sites.

|  | CNE7060-7061 | CNE7061-7063 | CNE6699-6700 | CNE6700-6697 | CNE6820-6816 | CNE6739-6741 |
| --- | --- | --- | --- | --- | --- | --- |
| Model selected | TVM+G | HKY85+I | HKY85 | TrN | HKY85 | K80+G |
| -lnL | 2716.1721 | 1036.8154 | 1035.1306 | 1182.1635 | 866.8431 | 730.5693 |
| Number of parameters | 8 | 5 | 4 | 5 | 4 | 2 |
| Base frequencies: |  |  |  |  |  |  |
| freqA = | 0.2998 | 0.2296 | 0.3066 | 0.2555 | 0.2128 | Equal frequencies |
| freqC = | 0.1780 | 0.1283 | 0.1908 | 0.2807 | 0.2338 |  |
| freqG = | 0.1813 | 0.1833 | 0.1764 | 0.1441 | 0.3864 |  |
| freqT = | 0.3409 | 0.4588 | 0.3263 | 0.3197 | 0.1670 |  |
| Substitution model: |  |  |  |  |  |  |
| Ti/tv ratio | Rate matrix | 0.9745 | 1.516 | Rate matrix | 1.9377 | 1.4804 |
| R(a) [A-C] | 2.7226 |  |  | 1 |  |  |
| R(b) [A-G] | 3.5734 |  |  | 6.1331 |  |  |
| R(c) [A-T] | 1.3229 | n/a | n/a | 1 | n/a | n/a |
| R(d) [C-G] | 0.6146 |  |  | 1 |  |  |
| R(e) [C-T] | 3.5734 |  |  | 2.9794 |  |  |
| R(f) [G-T] | 1 |  |  | 1 |  |  |
| Among-site rate variation: |  |  |  |  |  |  |
| Proportion of invariable sites | 0 | 0.3797 | 0 | 0 | 0 | 0 |
| Variable sites |  |  |  |  |  |  |
| Gamma distribution shape parameter $\alpha$ | 1.0216 | Equal rates for all sites | Equal rates for all sites | Equal rates for all sites | Equal rates for all sites | 0.243 |

The pattern of evolution of CNE6700-6697 can be described by the TrN model (Tamura and Nei 1993), which includes different nucleotide frequencies, one parameter for transitions and two for transversions. Furthermore, when arctic charr clones AC14, AC16, AC20 and AC21 are removed from the analysis, this locus’s evolution can be described according to a simpler model, HKY85 (see above) plus a gamma distribution of variable sites. In CNE7060-7061, the model chosen to explain how this locus evolves is the transversional model plus a gamma distribution of variable sites (TVM+G), including different nucleotide frequencies, one parameter for transitions and two for transversions. However, a different and simpler model, HKY85, describes the evolution of CNE7061-7063. In this case, another parameter is added, I, the proportion of sites that do not vary.

The homoplasy and rescaled consistency indexes estimate the data sets’ amount of phylogenetic information in the MP analyses. This last index ranged from 0.5704 to 0.9248 (Table 5) and is the ratio of the minimum number of steps required to build a tree with a given number of taxa and the actual number of steps in the tree (consistency index), corrected for the dataset size, as the consistency index is known to be sensitive to the number of taxa in the tree. Conversely, the amount of noise in the data, or the proportion of characters in a dataset that supports a different topology represented by the homoplasy index, ranges from 0.2667 to 0.463 (Table 5).

**Table 5.** Descriptive statistics for the Maximum Parsimony analyses of CNE clones from the three species of Salmonines, rainbow trout (*Oncorhynchus mykiss*), Atlantic salmon (*Salmo salar*) and Arctic charr (*Salvelinus alpinus*) included in this study.

|  | CNE7060-7061 | CNE7061-7063 | CNE6699-6700 | CNE6700-6697 | CNE6820-6816 | CNE6739-6741 |
| --- | --- | --- | --- | --- | --- | --- |
| Length (bp) | 676 | 252 | 357 | 539 | 289 | 265 |
| Variable sites | 267 | 102 | 88 | 78 | 89 | 49 |
| Parsimony-<br>informative sites | 144 | 66 | 44 | 58 | 57 | 29 |
| Tree length (steps) | 349 | 149 | 107 | 89 | 108 | 75 |
| Homoplasy index of<br>MP trees | 0.0926 | 0.1477 | 0.1028 | 0.0674 | 0.0463 | 0.2667 |
| Rescaled consistency<br>index | 0.8534 | 0.7974 | 0.7995 | 0.8962 | 0.9248 | 0.5704 |

### Phylogenetic analyses

Bayesian phylogenetic analyses show three of the six loci included in this study, namely CNE7060-7061, CNE7061-7063 and CNE6820-6816 (Figure 4*a*, 4*b* and 4*e* respectively), have a basal bifurcation with strong statistical support. Each of the resulting clades is comprised of clones from the three Salmoninae species. In these clades, the relationships between these three species are not resolved or are not resolved consistently across loci. In turn, these clades are not resolved at the species level, and two or more clones of the same species tend to stem from a single node (polytomies). On the other hand, loci CNE6699-6700, CNE6700-6697 and CNE6739-6741 are characterized by basal polytomies composed of many clades lacking statistical support. Overall, ML and MP analyses resulted in similar or identical topologies, which are highly consistent with those obtained in Bayesian analyses.

**Figure 4.**
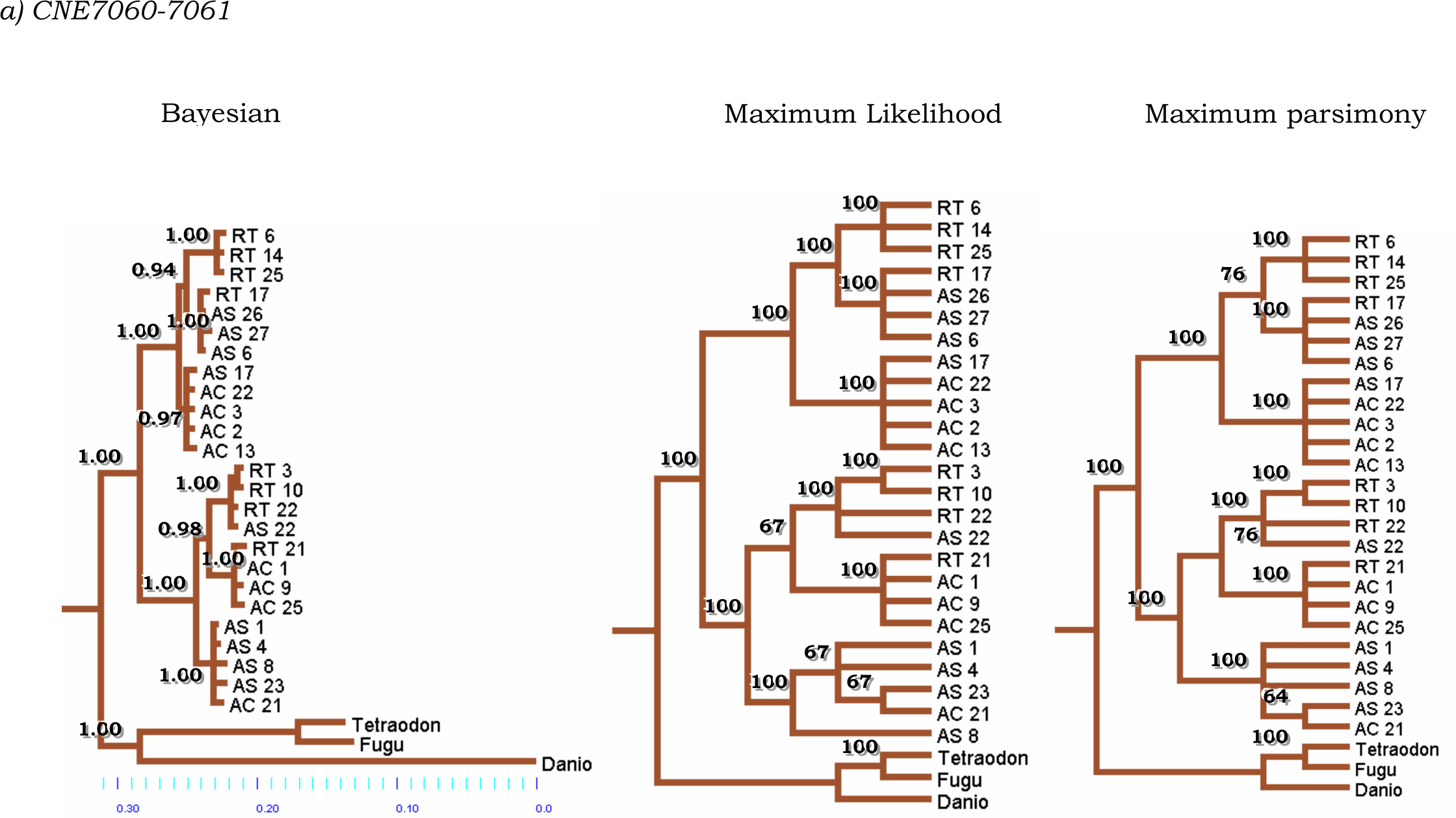
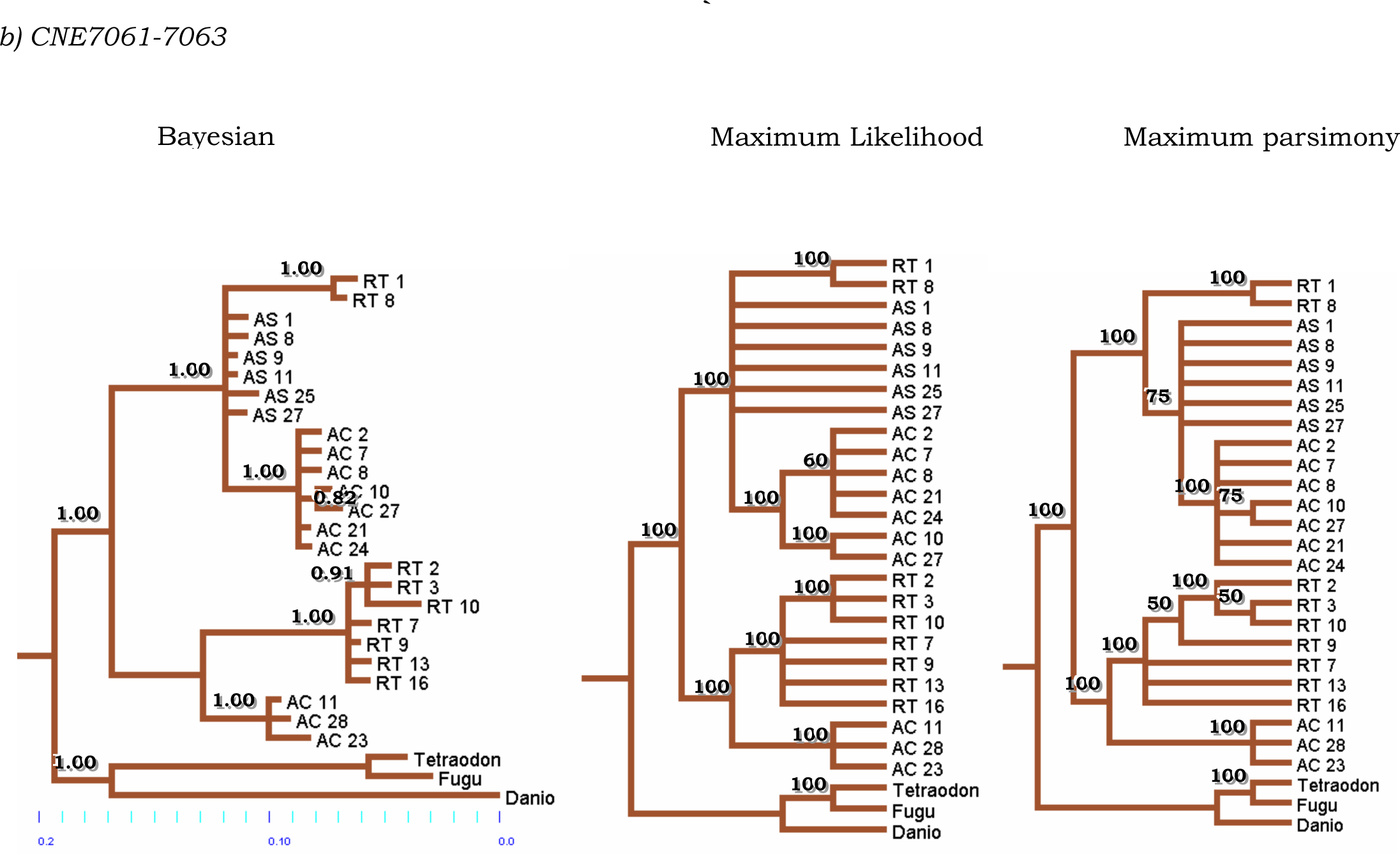
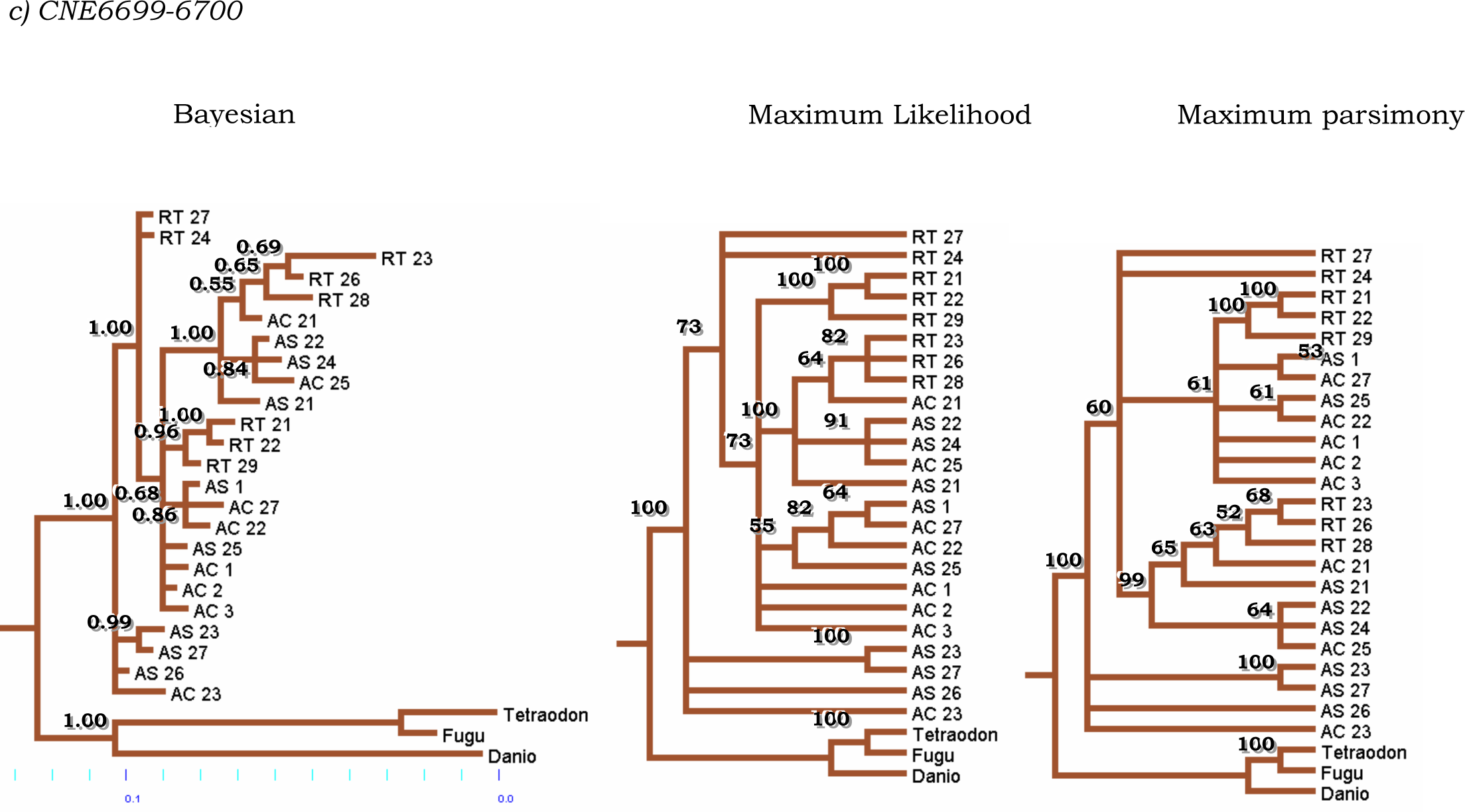
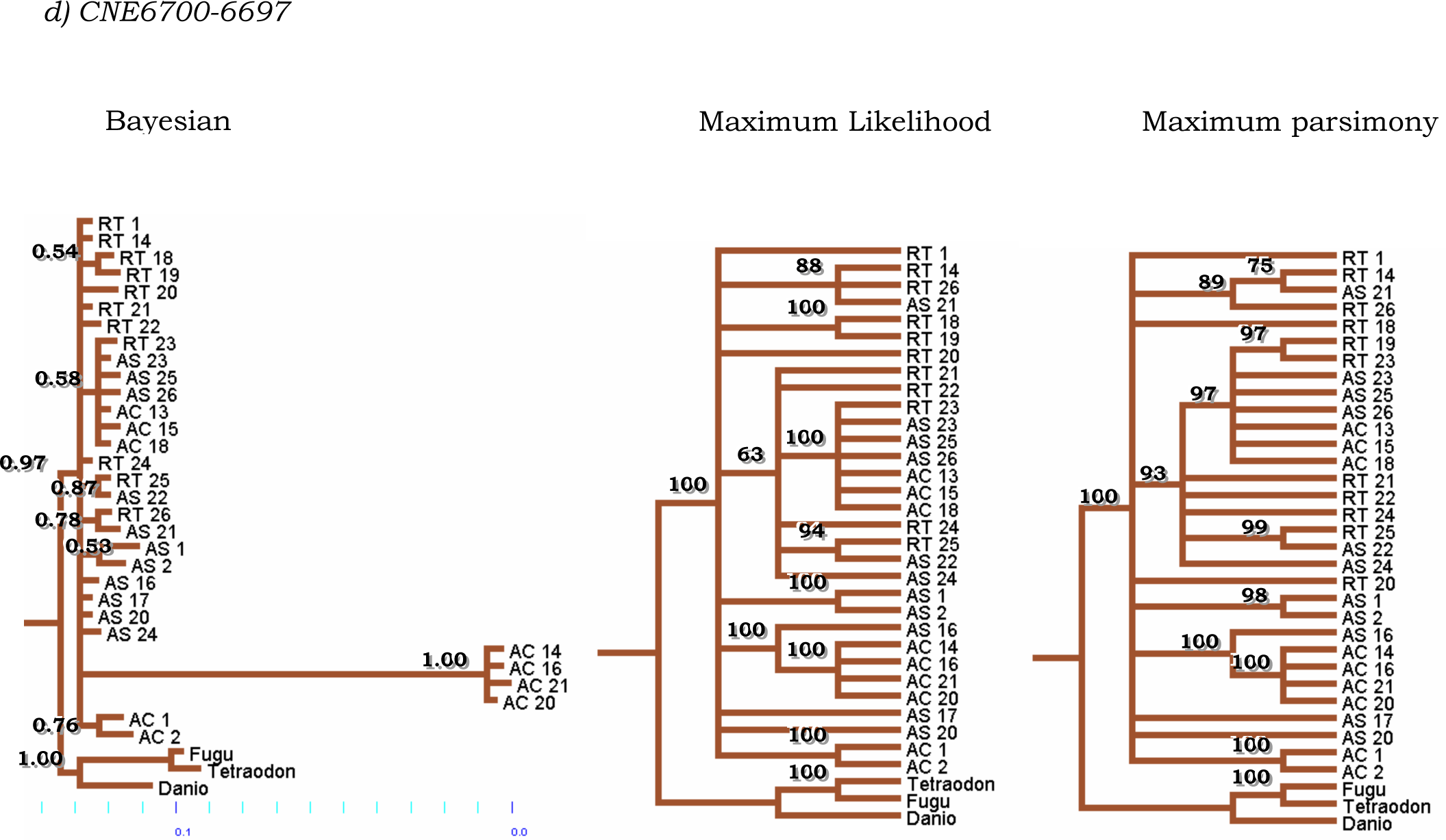
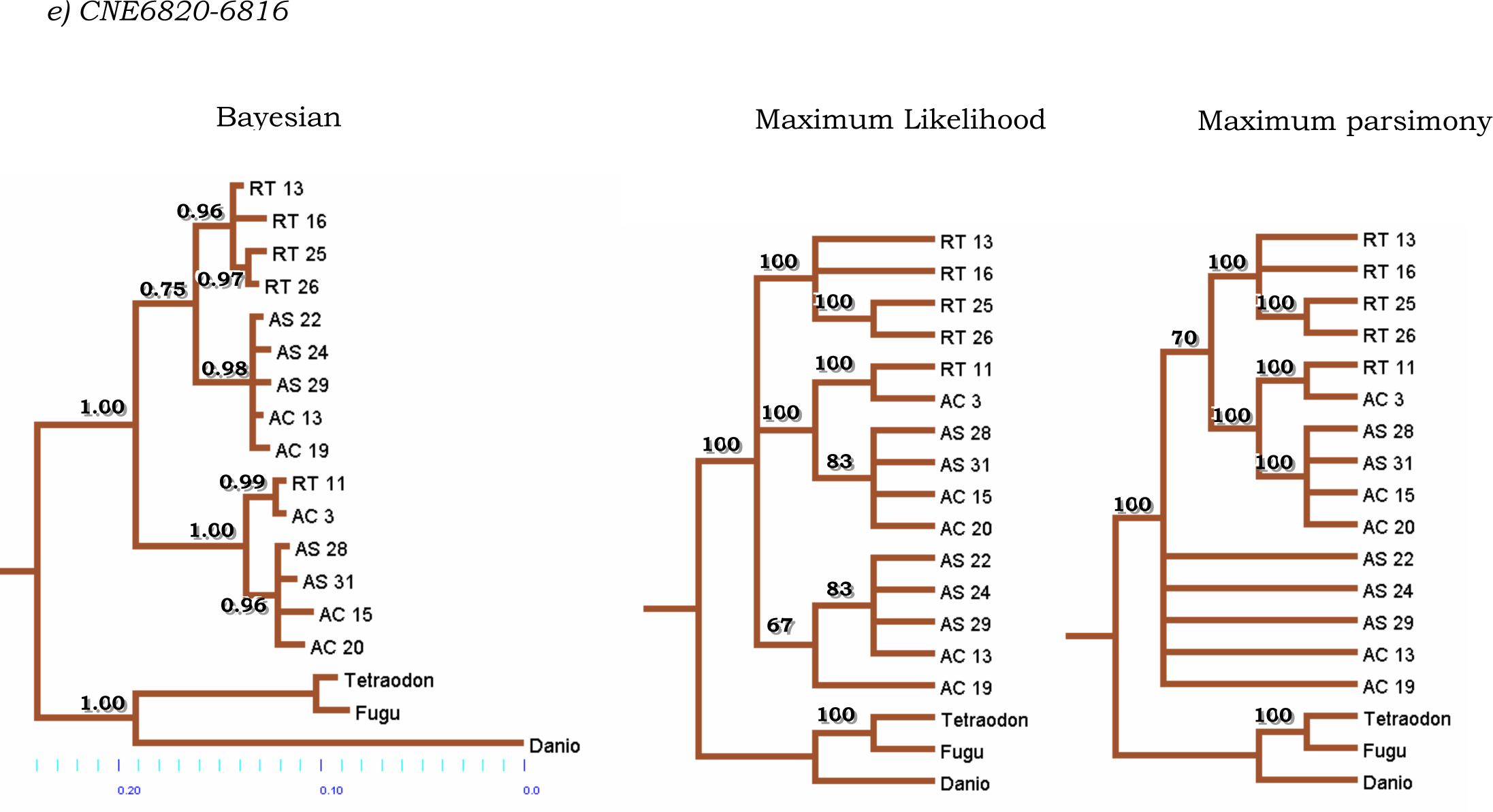
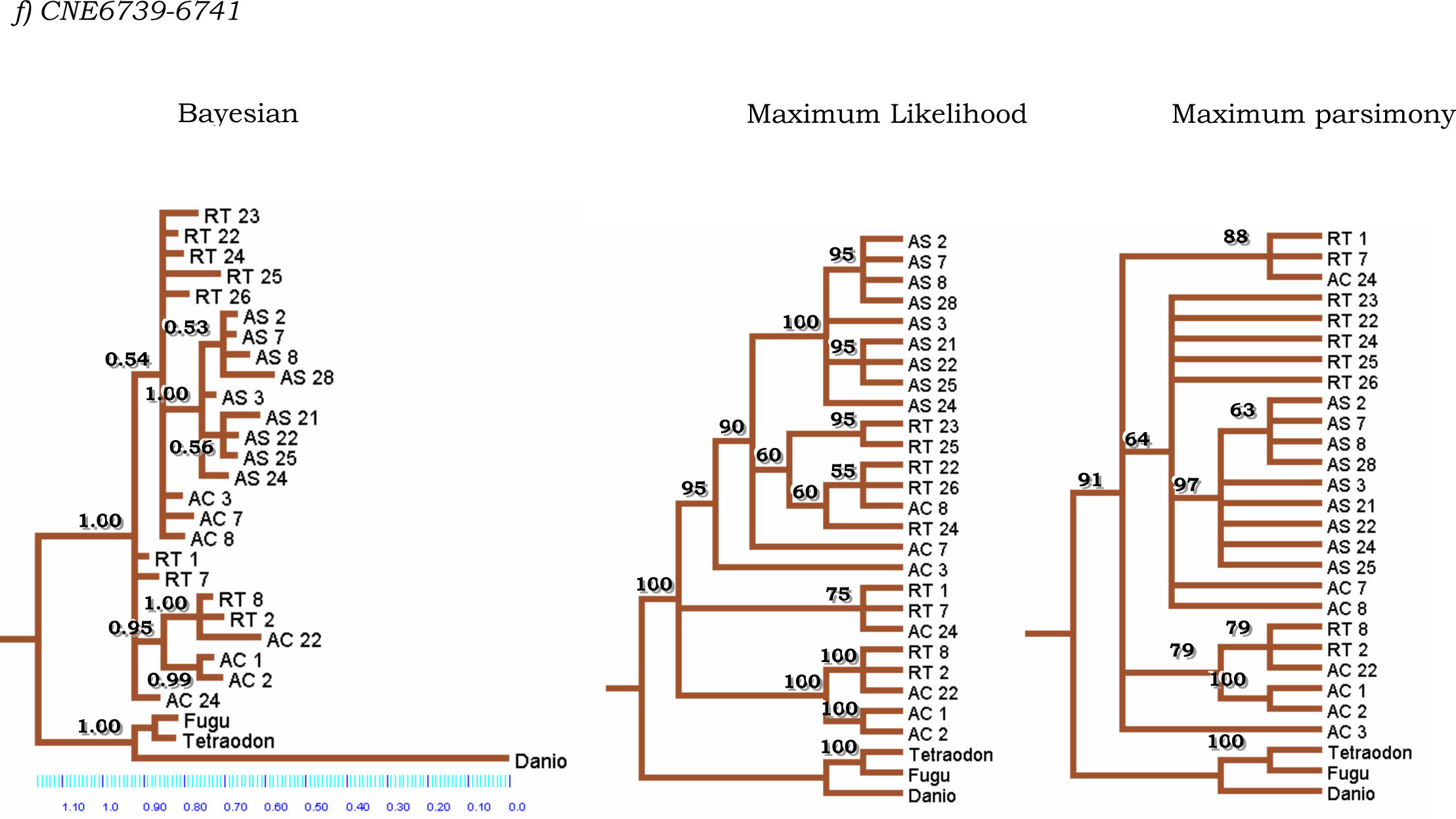
Phylogenetic relationships of CNE clones from three species of Salmonines, rainbow trout (*Oncorhynchus mykiss*), Atlantic salmon (*Salmo salar*) and Arctic charr (*Salvelinus alpinus*). The first tree corresponds to a Bayesian analysis. Numbers above branches indicate Bayesian posterior probabilities. The second tree corresponds to a consensus of the best trees obtained by Maximum Likelihood. Numbers above branches indicate consensus percentages. The third tree corresponds to a consensus of the most parsimonious trees. Numbers above branches indicate consensus percentages. — See ‘‘Materials and Methods’’ for details).

According to the Bayesian analyses, CNE7060-7061 shows a basal bifurcation resulting in two main clades (Figure 4*a*), both including representatives of the three species of Salmoninae. Each of these main clades, in turn, comprises smaller clades that are mostly composed of the three Salmoninae species as well. Interestingly, all these clades include one clone representative of another species. For example, a clade comprised of arctic charr clones AC2, AC3, AC13, AC22 (Figure 4*a*) also includes Atlantic salmon clone AS17. ML and MP analyses yielded the same phylogeny, except that clones AS23 and AC21 cluster together, and they, in turn, cluster with AS4 and AS1. In the BA, all of them collapse into a polytomy with AS8.

Moreover, locus CNE7061-7063 also shows a basal bifurcation resulting in two main clades (Figure 4*b*). One clade includes clones from the three Salmoninae species, with two rainbow trout clones (RT1 and RT8) grouping together, and a clade comprising seven arctic charr clones. These two groups, together with six Atlantic salmon clones, form a polytomy. The other main basal clade includes only rainbow trout and arctic charr clones, both constituting polytomies. The clades mentioned above bear a high level of statistical support, as indicated by their posterior probabilities. It is worth noticing that ML and MP analyses yielded the same results, also with strong support. ML differs in that AC2, AC7, AC8, AC21, and AC24 form a clade together instead of a polytomy with the clade AC10-AC27. In turn, MP differs from BA in the level of resolution of the clade (RT2-RT3-RT7-RT9-RT10-RT13-RT16) (compare Bayesian and ML trees).

In contrast, locus CNE6699-6700 shows a polytomy near the root of the tree (basal), from which one main clade stems (Figure 4*c*). This clade lacks strong support and, in turn, is itself another polytomy that includes one highly supported clade of Salmoninae. ML and MP yielded similar results. Likewise, the phylogeny of locus CNE6700-6697 is characterized by a main basal polytomy (Figure 4*d*). It includes one poorly supported clade with clones from the three species of Salmoninae and poorly supported groups comprised of one pair of rainbow trout clones and one of Atlantic salmon clones.

Moreover, stemming from this basal polytomy are two clades comprised of one rainbow trout clone and one Atlantic salmon clone. Noticeably, there is one clade formed by four arctic charr clones that stems from the polytomy through a very long branch. Coincidentally, these clones significantly differ in length from the rest of the clones, even those that also belong to arctic charr (Table 3). Once again, the results reported by ML and MP are highly consistent with those of the BA.

The phylogeny of locus CNE6820-6816 (Figure 4*e*) shows a basal bifurcation. The first one is further divided into two groups: rainbow trout clones and one consisting of both arctic charr and Atlantic salmon clones. The second main clade consists of two clades, one comprised of an arctic charr clone and a rainbow trout clone, and the other one including arctic charr and Atlantic salmon clones. All the groups mentioned above bear substantial statistical support. ML yielded the same phylogeny and according to MP, the second clade mentioned above collapses into a polytomy.

Similar to CNE6699-6700, CNE6739-6741 shows a basal polytomy (Figure 4*f*), from which two main groups originate: one highly supported clade of rainbow trout and arctic charr clones and a poorly supported polytomy that includes a well-supported clade of Atlantic salmon clones among rainbow trout and arctic charr clones. This result is consistent with ML and MP analyses.

### Molecular clock

The molecular clock hypothesis was rejected for all loci except CNE6820-6816 (Table 6). According to the molecular clock, dating the duplication event in this locus (see Figure 4*e* for a detailed phylogeny) resulted in an estimate of at least 47My with a 95% confidence interval of 32 - 65My (Figure 5). Conversely, locus CNE7060-7061 (Figure 4*a*) resulted in an estimate of at least 38My with a 95% confidence interval of 23 - 60My and locus CNE7061-7063 (Figure 4*b*), in turn, suggests the duplication event occurred at least 43Mya with a 95% confidence interval of 22 - 68Mya.

**Figure 5.**
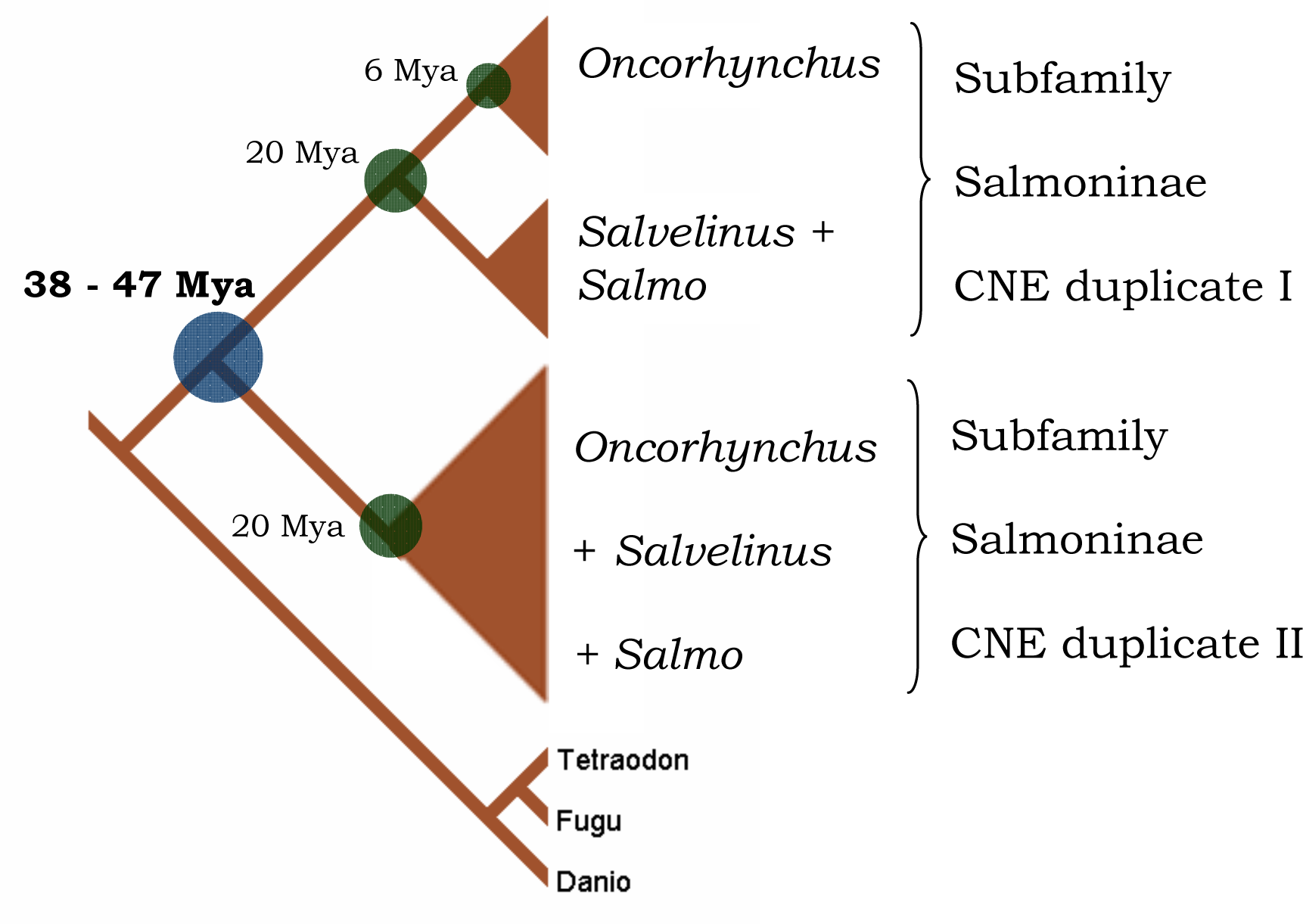
Dating of the duplication event in CNE6820-6816, CNE7060-7061 and CNE7061-7063 according to the molecular clock hypothesis (see figures 4a, b and e for detailed phylogenies). The following calibration points were used: 1) nodes including only rainbow trout clones were considered representatives of the origin of the genus *Oncorhynchus* and were therefore assigned an age of six million years (My) according to Nelson (2006, p. 202); 2) clades including the genera *Salmo*, *Salvelinus* and *Oncorhynchus* were considered representatives of the origin of the subfamily Salmoninae and were assigned an age of 20My according to Groot (1996). This resulted in an estimate of at least 38-47My for the 4R genome duplication.

**Table 6.** Test of the molecular clock hypothesis for each pair of CNE. Indicated are the model used, the ln-likelihood of the model enforcing and not enforcing a molecular clock and the value of the Likelihood Ratio Test with its corresponding degrees of freedom and critical values.

| Locus | model | -ln L0 clock | -ln L1 no clock | LRT* | df** | critical value (P = 0.05) | Molecular clock hypothesis |
| --- | --- | --- | --- | --- | --- | --- | --- |
| CNE7060-7061 | HKY+G | 2752.4755 | 2727.2290 | 50.4930 | 26 | 38.89 | *Rejected* |
| CNE7061-7063 | HKY+G | 1076.7494 | 1007.7713 | 137.9562 | 26 | 38.89 | *Rejected* |
| CNE6699-6700 | HKY | 1065.7567 | 1035.1306 | 61.2522 | 25 | 37.65 | *Rejected* |
| CNE6700-6697 | TrN | 1253.4033 | 1182.1635 | 142.4797 | 32 | 46.19 | *Rejected* |
| CNE6820-6816 | HKY | 875.9860 | 866.8436 | 18.2848 | 16 | 26.30 | *Accepted* |
| CNE6739-6741 | K80+G | 756.4716 | 730.5693 | 51.8047 | 26 | 38.89 | *Rejected* |

\* LRT = 2 x (lnL<sub>1</sub>-lnL<sub>0</sub>)
\*\* df (degrees of freedom) is equivalent to the number of taxa minus two (Felsenstein 1981)

## Discussion

The number of different copies and nucleotide variation in six pairs of CNEs provides evidence that they follow the pattern expected based on Salmonids’ 4R duplication. Locus CNE6699-6700 and CNE6739-6741, which appear as double copies in zebrafish, showed about eight different alleles for each species, as predicted.

In some cases, more sequences were detected than predicted. This observation could be explained by the presence of more than twice as many loci as there are in zebrafish, *Taq* polymerase incorporation, or sequencing errors. Five alleles were detected in each species for locus CNE6820-6816, which has been retained as a single copy in zebrafish. Similarly, loci CNE7060-7061 and CNE7061-7063, which occur as single copies in zebrafish, show more than twice as many alleles in all three species. Arctic charr, however, shows two different groups of sequences for CNE6700-6697, including four and five sequences, respectively, which could be interpreted as belonging to at least two different loci. It is possible, although unlikely that further annotation of the zebrafish genome in the Ensembl database will reveal another copy of these loci. In turn, numbers of sequences lower than expected could be explained by the individuals from which these sequences were obtained being homozygotes for these loci, although this possibility should be confirmed.

Basal bifurcations in the phylogenetic trees for CNE7060-7061, CNE7061-7063 and CNE6820-6816 also suggest that these CNEs follow the expected pattern based on the 4R hypothesis (Figure 4*a*, *b* and *e*, respectively). The fact that each of these CNE pairs generated trees with two main clades with strong statistical support that include all three species supports the idea that the two clades constitute different, duplicated loci (Friedman and Hughes 2001; Steinke et al. 2006). Similar results were found by Moghadam et al. (2005), through an analysis of Hox genes’ genomic organization in Atlantic salmon and rainbow trout.

Interestingly, although CNE7061-7063 shows a basal bifurcation, its phylogeny is not symmetric, as one of the clades does not include Atlantic salmon. Phylogenetic signal might be lacking for this locus in this species, given that all its clones collapse into a basal polytomy. The topologies of the remaining loci, CNE 6697-6700, CNE6700-6697 and CNE6739-6741, constitute polytomies that do not contribute to the resolution of this phylogenetic conundrum and therefore do not conform to the expected patterns resulting from the 4R genome duplication. The phylogenetic signal might be lacking for these loci in these species, or the level of nucleotide variation might not be appropriate for the time frame and taxonomic level (millions of years and genera) being addressed.

The six pairs of CNE loci were of value in dating the 4R genome duplication. Three CNE pairs, CNE7060-7061, CNE7061-7063 and CNE6820-6816, could achieve this goal because they showed a basal bifurcation (see above). The estimated dates for the lower limit of the 4R duplication of the Salmonine genome, ranging between 38 to 47My, are consistent with the reported time range of a putative WGD in the Salmonids ancestor 25-100Mya (Allendorf and Thorgaard1984). It is noteworthy that the present work estimates are the first for the 4R WGD in the Salmonine fishes’ ancestor since the pioneering work of Allendorf and Thorgaard (1984) and the first-ever based on nucleotide sequence data. Therefore, this is an update on an issue that awaited further refinement for almost 25 years (Gregory and Mable 2005).

Nevertheless, as is usual with calibrations of the molecular clock, many factors render this estimate uncertain. For instance, the stochastic nature of the molecular clock (Hillis et al. 1996; Benton and Ayala 2003), phylogenetic error, the uncertainty of the assigned fossil dates and even the uncertainty of the correspondence between the fossils reported in the literature and the nodes I assumed they represent. Furthermore, this attempt to date a duplication in Salmonidae’s ancestor only included representatives of one subfamily, Salmoninae. Other subfamilies within Salmonidae should be included (e.g., Thymallinae) to obtain more robust results. Therefore, this may be considered a narrower, but still rough estimate of the 4R genome duplication date.

It is worth mentioning that in CNE6700-6697, arctic charr clones AC16, AC16, AC20 and AC21 have a set of insertions compared to other clones from arctic charr and clones from rainbow trout and Atlantic salmon. One of these insertions occurs inside CNE CR846697 (see Appendix I). This result is noteworthy, as this CNE is 92.2% similar when compared between Japanese pufferfish and humans (Woolfe et al. 2005). It has been shown that some CNEs are involved in the regulation of early development genes in vertebrates (Woolfe et al. 2005).

However, since the exact role CNEs is not yet known, it is not possible to deduce, without functional analyses, if these longer arctic charr alleles are functional or not. Moreover, this could be a case of subfunctionalization arising from a duplication (Lynch and Force 2000a,b).

More than a single duplication is needed to explain the evolution of loci CNE7060-7061, CNE7061-7063 and CNE6820-6816. If one duplication accounted for these loci’s evolution, every clade in their phylogenetic trees would only have up to two alleles. In all cases, more alleles were found (Table 3). If there had been another WGD or segmental duplication of these loci in the ancestor of Salmonines, then the tree topology of these loci would be different from the one shown in Figures 4*a*, *b* and *e*. Each of the main clades would be split into two clades that would include the three species of Salmonines, containing up to two alleles in each clade.

On the contrary, the scenario that best explains the number of alleles and their phylogenetic topology is a segmental duplication in each species following the 4R genome duplication. As mentioned before, such scenarios have been considered by Robinson-Rechavi et al. (2001a,d), who suggested that the number of copies in some gene families in teleost fishes is best explained by segmental duplications rather than genome duplication (see also Taylor et al. 2001c and Robinson-Rechavi et al. 2001b). Moreover, Kawasaki et al. (2007) suggest that the secretory calcium-binding phosphoprotein gene family, responsible in part for the mineralized skeleton in vertebrates, owes its genetic architecture to a combination of genome and gene duplications.

It is unclear if the basal polytomies shown in the present study could be interpreted as the result of an adaptive radiation, as there is not much information about the conditions that could have led to such process 20Mya, the estimated age of the ancestor of the subfamily Salmoninae (Groot 1996). However, it has been suggested that the tetraploidization event that Salmonids underwent between 25 to 100Mya has played a significant role in promoting their diversification (Allendorf and Thorgaard 1984). In this regard, it has also been suggested that polyploidization events coincide with significant taxonomic diversification in ray-finned lineages and flowering plants (Van de Peer and Meyer 2005 and references therein). Van de Peer and Meyer (2005) suggest that the family Salmonidae has significantly more species than the related group Esocidae might be associated with divergent resolution (differential loss of duplicates) following tetraploidization. Kinnison and Hendry (2004) suggest that the fact that said adaptive radiation had not been mirrored by the also tetraploid related subfamily Thymallinae is an indication that factors other than the tetraploidization event must have played a role in the diversification of Salmonids. There are documented cases where sister taxa do not undergo the same speciation rate. For instance, the rodent genus *Ctenomys* has undergone an increased rate of diversification compared to their sister taxon, the octodontids (Castillo et al. 2005) and the reasons for this differential speciation rate are probably found at the ecological and population genetics level (see Castillo et al. 2005 and references therein).

### CNEs as phylogenetic tools

The phylogenies for the six pairs of CNEs did little to resolve the phylogenetic affinities of the three genera of Salmoninae, as each locus that showed phylogenetic resolution at the species level presented a different result. The two duplicates in CNE7060-7061 show different results. One of the duplicates shows Atlantic salmon as the sister group of rainbow trout (main clade at the top of Figure 4*a*). Similar results have been previously reported (Phillips and Pleyte 1991; Stearley and Smith 1993; Murata et al. 1996; Phillips and Oakley 1997). In contrast, the second duplicate in CNE7060-7061 shows rainbow trout as being more related to arctic charr, coinciding with the findings of Oakley and Phillips (1999) and Crespi and Fulton (2003), who suggested that the genera *Oncorhynchus* and *Salvelinus* are more closely related to each other than either of them is to *Salmo*. A similar result is shown by CNE6739-6741, although many clones in the tree collapse into a polytomy. Interestingly, both duplicates in CNE6820-6816 (Figure 4*e*) show Atlantic salmon and arctic charr being more closely related to each other than either of them is to rainbow trout. This relationship has not been previously reported.

Moreover, it is known that gene trees and species trees might differ (Maddison 1997; Nakhleh et al. 2005). The rest of the CNE pairs did not show phylogenetic resolution at the species level. Possible reasons for this result include reticulate evolution and the fact that these specific loci occur as duplicate copies in zebrafish. Given that, in turn, they are duplicated in Salmonines, it is expected that this would distort the phylogenetic signal, and the timing of the speciation events within the subfamily Salmoninae is beyond the temporal resolution of these loci.

Given the strong support of these phylogenies indicated by the posterior probabilities and the high rescaled consistency indexes, it is unlikely that this inconsistency across loci is due to lack of phylogenetic signal in the data. This last case is known as a “soft polytomy” (Maddison and Maddison 1992; Coddington and Scharff 1996). When any event or process is tested by tree topology, time must have passed since the event/process took place so that a phylogenetic signal can reveal it. Therefore, a soft polytomy can be thought of as branching of lineages being fast enough that the internode length is short and lacks statistical support (see Hrbeck et al. 2005 for a discussion of the fish genus *Aphanius*).

Alternatively, these loci’s inability to resolve the Salmonines’ phylogeny might be due to the separation of the three genera having co-occurred in geological time, which is known as a “hard polytomy” (Maddison and Maddison 1992; Coddington and Scharff 1996; Walsh et al. 1999), and can result from rapid speciation events and adaptive radiations. These processes have been documented in many taxa and particularly in fishes, one of the most typical cases being the adaptive radiation of cichlids (Seehausen 2006 and references therein). Furthermore, adaptive radiations have been invoked to explain hard polytomies in the fish genus *Chondrostoma* (Durand et al. 2003, but see also Hrbeck et al. 2002 for a discussion of the case of the fish genus *Aphanius*).

From the data collected in this study regarding the number of alleles per locus, tree topology and the estimated date for a potential genome duplication, there is partial evidence that CNEs can recover the 4R genome duplication in Salmonines. More robust conclusions could be drawn by investigating more CNE loci, collecting more sequence data within individual fish species to estimate correct allele copy numbers, and including more salmonid species. These results would be complemented by mapping different copies of a given CNE on different LG in Salmonines, to help identify pairs of known homeologues and relate the two pairs as being derived from one of the twelve ancestral proto-vertebrate LG (*sensu* Jaillon et al. 2004). If the two duplicates map to known homeologue chromosomes in Salmonines, it would indicate a WGD. If, in turn, they map to the same chromosome, it would be suggestive of a tandem repeat. Attempts were made in the present study to gather this information but were unsuccessful, as, at the most, only one duplicate could be mapped at a time (results not shown). This information helps improve the density of these species’ genetic maps but does not help discern between genomic evolution scenarios mentioned above.

Much of the knowledge gained about the genetic architecture in Salmonine species comes from comparative analyses of available genetic maps for different species (Danzmann et al. 2005). Even in taxa of longstanding interest like the amphibian genus *Xenopus,* where different levels of polyploidy have been reported (Evans et al. 2004), genome sequencing and mapping efforts are underway only for the model species *X. tropicalis*, so it is unlikely that comparative analyses will be available soon. Nevertheless, the history of polyploidization events in this genus has been successfully addressed through a phylogenetic approach (Evan et al. 2004). CNEs can be considered a suitable molecular tool to investigate genome duplications before more comprehensive and demanding endeavours are carried out.

## Acknowledgements

Roy G. Danzmann, Hooman Moghadam and Moira M. Ferguson provided valuable advice throughout this study and helpful comments and discussions on earlier drafts of this manuscript and throughout the writing process.

## References

Akaike, H. 1974. A new look at the statistical model identification. IEEE Transactions on Automatic Control 19 (6): 716–723

Allendorf F.W. and G.H. Thorgaard. 1984. Tetraploidy and the evolution of salmonid fishes. In: Turner JB (Ed) Evolutionary genetics of fishes. Plenum Press, New York, pp 1–53

Benton, M.J. and F.J. Ayala. 2003. Dating the tree of life. Science 300: 1698–1700

Bos, D.H. and D. Posada. 2005. Using models of nucleotide evolution to build phylogenetic trees. Developmental and Comparative Immunology 29: 211–227

Castillo, A.H., M.N. Cortinas, and E.P. Lessa. 2005. Rapid diversification of South American tuco-tucos (*Ctenomys*; Rodentia, Ctenomyidae): Contrasting mitochondrial and nuclear intron sequences. Journal of Mammalogy 86: 170–179

Coddington, J.A. and N. Scharff. 1996. Problems with “soft” polytomies. Cladistics-the International Journal of the Willi Hennig Society 12: 139–145

Crespi, B.J. and M.J. Fulton. 2004. Molecular systematics of Salmonidae: combined nuclear data yields a robust phylogeny. Molecular Phylogenetics and Evolution 31: 658–679

Danzmann R.G., Cairney M., Davidson W.S., Ferguson M.M., Gharbi K., Guyomard R., Holm L.E., Leder E., Okamoto N., Ozaki A., Rexroad C.E., Sakamoto T., Taggart J.B. and R.A. Woram. 2005. A comparative analysis of the rainbow trout genome with two other species of fish (Arctic charr and Atlantic salmon) within the tetraploid derivative Salmonidae family (subfamily: Salmoninae). Genome 48: 1037–1051

Drummond, A.J., SYW Ho, M.J. Phillips, and A. Rambaut. 2006. Relaxed phylogenetics and dating with confidence. PLoS Biology 4: 699–710

Durand, J.D., P.G. Bianco, J. Laroche, and A. Gilles. 2003. Insight into the origin of endemic Mediterranean ichthyofauna: Phylogeography of *Chondrostoma* genus (Teleostei, Cyprinidae). Journal of Heredity 94: 315–328

Eichler E.E. and NH. Patel. 2003. Genomes and evolution: From sequence to organism. Current Opinion in Genetics and Development 13: 559–561

Evans, B.J., D.B. Kelley, R.C. Tinsley, D.J. Melnick, and D.C. Cannatella. 2004. A mitochondrial DNA phylogeny of African clawed frogs: phylogeography and implications for polyploid evolution. Molecular Phylogenetics and Evolution 33: 197–213

Felsenstein, J. 1981. Evolutionary trees from DNA sequences: a maximum likelihood approach. Journal of Molecular Evolution 17: 368–376

Froschauer, A., I. Braasch, and J.N. Volff. 2006. Fish genomes, comparative genomics and vertebrate evolution. Current Genomics 7: 43–57

Friedman, R. and A.L. Hughes. 2001. Pattern and timing of gene duplication in animal genomes. Genome Research 11: 1842–1847

Furlong, R.F. and PWH Holland. 2002. Were vertebrates octoploid? Philosophical Transactions of the Royal Society of London Series B-Biological Sciences 357: 531–544

Gallardo, M.H., J.W. Bickham, R.L. Honeycutt, R.A. Ojeda, and N. Kohler. 1999. Discovery of tetraploidy in a mammal. Nature 401: 341–341

Gallardo, M.H., J.W. Bickham, G. Kausel, N. Kohler, and R.L. Honeycutt. 2003. Gradual and quantum genome size shifts in the hystricognath rodents. Journal of Evolutionary Biology 16: 163–169

Gallardo, M.H., C.A. Gonzalez, and I. Cebrian. 2006. Molecular cytogenetics and allotetraploidy in the red vizcacha rat, Tympanoctomys barrerae (Rodentia, Octodontidae). Genomics 88: 214–221

Gallardo, M.H., G. Kausel, A. Jimenez, C. Bacquet, C. Gonzalez, J. Figueroa, N. Kohler, and R. Ojeda. 2004. Whole-genome duplications in South American desert rodents (Octodontidae). Biological Journal of the Linnean Society 82: 443–451

Gregory T.R. (editor) (2005). The Evolution of the Genome. Elsevier, San Diego.

Gregory T.R. and BK. Mable (2005). Polyploidy in animals. In: The Evolution of the Genome, edited by T.R. Gregory. Elsevier, San Diego, pp. 427–517

Groot C. 1996. Principles of salmonid culture. In: W. Pennell and BA. Barton [Eds.]. Salmonid Life Histories. Elsevier. Developments in Aquaculture and Fisheries Science, 29: 97–230

Hasegawa M., Kishino K. and T. Yano 1985. Dating the human-ape splitting by a molecular clock of mitochondrial DNA. Journal of Molecular Evolution 22:160–174

Hubbard T. J. P., B. L. Aken, K. Beal, B. Ballester, M. Caccamo, Y. Chen, L. Clarke, G. Coates, F. Cunningham, T. Cutts, T. Down, S. C. Dyer, S. Fitzgerald, J. Fernandez-Banet, S. Graf, S. Haider, M. Hammond, J. Herrero, R. Holland, K. Howe, K. Howe, N. Johnson, A. Kahari, D. Keefe, F. Kokocinski, E. Kulesha, D. Lawson, I. Longden, C. Melsopp, K. Megy, P. Meidl, B. Overduin, A. Parker, A. Prlic, S. Rice, D. Rios, M. Schuster, I. Sealy, J. Severin, G. Slater, D. Smedley, G. Spudich, S. Trevanion, A. Vilella, J. Vogel, S. White, M. Wood, T. Cox, Curwen, R. Durbin, X. M. Fernandez-Suarez, P. Flicek, A. Kasprzyk, G. Proctor, S. Searle, J. Smith, A. Ureta-Vidal and E. Birney. 2007. Ensembl 2007. Nucleic Acids Research 35: D610–D617

Huelsenbeck, JP and F. Ronquist. 2001. MrBayes: Bayesian inference of phylogeny. Bioinformatics 17: 754–755

Jackson, T.R., M.M. Ferguson, R.G. Danzmann, A.G. Fishback, P.E. Ihssen, M. O’Connell, and TJ Crease. 1998. Identification of two QTL influencing upper temperature tolerance in three rainbow trout (*Oncorhynchus mykiss*) half-sib families. Heredity 80: 143–151

Jaillon O., Aury J.M., Brunet F., Petit J.L., Stange-Thomann N., Mauceli E., Bouneau L., Fischer C., Ozouf-Costaz C., Bernot A., Nicaud S., Jaffe D., Fisher S., Lutfalla G., Dossat C., Segurens B., Dasilva C., Salanoubat M., Levy M., Boudet N., Castellano S., Anthouard V., Jubin C., Castelli V., Katinlca M., Vagherie B., Biemont C., Skalli Z., Cattolico L., Poulain J., De Berardinis V., Cruaud C., Duprat S., Brottier P., Coutanceau J.P., Gouzy J., Parra G., Lardier G., Chapple C., McKernan K.J., McEwan P., Bosak S., Kellis M., Volff J.N., Guigo R., Zody M.C., Mesirov J., Lindblad-Toh K., Birren B., Nusbaum C., Kahn D., Robinson-Rechavi M., Laudet V., Schachter V., Quetier F., Saurin W., Scarpelli C., Wincker P., Lander E.S., Weissenbach J. and H. Roest Crollius. 2004. Genome duplication in the teleost fish *Tetraodon nigroviridis* reveals the early vertebrate proto-karyotype. Nature 431: 946–957

Jukes T.H. and C.R. Cantor. 1969. Evolution of protein molecules. In: Munro HM (Ed): Mammalian Protein Metabolism. Academic Press, New York, NY, p 21–132

Kasahara M. 2007. The 2R hypothesis: an update. Current Opinion in Immunology 19:547–552

Kawasaki, K., A.V. Buchanan, and K.M. Weiss. 2007. Gene duplication and the evolution of vertebrate skeletal mineralization. Cells Tissues Organs 186: 7–24

Kellogg, E.A. 2003a. It’s all relative. Nature 422: 383–384

Kellogg, E.A. 2003b. What happens to genes in duplicated genomes. Proceedings of the National Academy of Sciences of the United States of America 100: 4369–4371

Kimura M. 1980. A simple method for estimating evolutionary rate of base substitutions through comparative studies of nucleotide sequences. Journal of Molecular Evolution 16: 111–120

Kinnison, M.T. and A.P. Hendry. 2003. From macro- to micro-evolution - Tempo and mode in Salmonid evolution. In: A.P. Hendry (Ed) Evolution Illuminated: Salmon and Their Relatives, Oxford Press, New York, pp 208–231

Larhammar D., Lundin L.G. and F. Hallbook. 2002. The human Hox-bearing chromosome regions did arise by block or chromosome (or even genome) duplications. Genome Research 12: 1910–1920

Lewin R. 1987. National academy looks at human genome project, sees progress. Science 235: 747–748

Lio, P. and N. Goldman. 1998. Models of molecular evolution and phylogeny. Genome Research 8: 1233–1244

Lundin, L.G. 1993. Evolution of the vertebrate genome as reflected in paralogous chromosomal regions in man and the house mouse. Genomics 16: 1–19

Lynch, M. and A. Force. 2000a. The probability of duplicate gene preservation by subfunctionalization. Genetics 154: 459–473

Lynch, M. and A.G. Force. 2000b. The origin of interspecific genomic incompatibility via gene duplication. American Naturalist 156: 590–605

Maddison, W.P. 1997. Gene trees in species trees. Systematic Biology 46: 523–536

Maddison, W. P., and Maddison, D. R. 1992. MacClade, version 3.0. Sinauer, Sunderland, MA.

Massingham T., L.J. Davies, and P. Lio. 2001. Analysing gene function after duplication. Bioessays 23: 873–876

Mackay T.F.C. (2001). The genetic architecture of quantitative traits. Annual Review of Genetics 35: 303–339

Meyer A. and Y. Van de Peer. 2003. Genome Evolution: Gene and Genome Duplications and the Origin of Novel Gene Functions. Springer.

Moghadam H.K., Ferguson M.M. and R.G. Danzmann. 2005. Evolution of *Hox* clusters in Salmonidae: a comparative analysis between Atlantic salmon *(Salmo salar)* and rainbow trout *(Oncorhynchus mykiss)*. Journal of Molecular Evolution 61: 636–649

Murata, S., N. Takasaki, M. Saitoh, H. Tachida, and N. Okada. 1996. Details of retropositional genome dynamics that provide a rationale for a generic division: The distinct branching of all the Pacific salmon and trout (*Oncorhynchus*) from the Atlantic salmon and trout (*Salmo*). Genetics 142: 915–926

Nakhleh, L., T. Warnow, C.R. Linder, and K. St John. 2005. Reconstructing reticulate evolution in species - Theory and practice. Journal of Computational Biology 12: 796–811

Nylander, J. A. A. 2004. MrModeltest v2. Program distributed by the author. Evolutionary Biology Centre, Uppsala University. www.abc.se/~nylander

Oakley, T.H. and R.B. Phillips. 1999. Phylogeny of salmonine fishes based on growth hormone introns: Atlantic (*Salmo*) and Pacific (*Oncorhynchus*) salmon are not sister taxa. Molecular Phylogenetics and Evolution 11: 381–393

Ohno S. 1970. Evolution by gene duplication. Springer Verlag, New York

Ohno S. 1999. Gene duplication and the uniqueness of vertebrate genomes circa 1970-1999. Cell and Developmental Biology. 10: 517–522

Phillips R.B. and P. Ráb. 2001. Chromosome evolution in the Salmonidae (Pisces): an update. Biological Reviews 76: 1–25

Phillips R.B. and K.A. Pleyte. 1991. Nuclear-DNA and salmonid phylogenetics. Journal of Fish Biology 39: 259–275

Posada D. and K.A. Crandall. 1998. Modeltest: testing the model of DNA substitution. Bioinformatics 14: 817–818

Posada, D. and K.A. Crandall. 2001a. Selecting the best-fit model of nucleotide substitution. Systematic Biology 50: 580–601

Posada, D. and K.A. Crandall. 2001b. Simple (wrong) models for complex trees: A case from retroviridae. Molecular Biology and Evolution 18: 271–275

Posada, D. and T.R. Buckley. 2004. Model selection and model averaging in phylogenetics: Advantages of Akaike information criterion and Bayesian approaches over likelihood ratio tests. Systematic Biology 53: 793–808

Rambaut A. and A.J. Drummond. 2007 Tracer v1.4, Available from http://beast.bio.ed.ac.uk/Tracer

Robinson-Rechavi M., Marchand O., Escriva H. and V. Laudet. 2001a. Ancestral whole-genome duplications may not have been responsible for abundance of duplicated fish genes. Current Biology 11: R458–R459

Robinson-Rechavi M., Marchand O., Escriva H. and V. Laudet. 2001b. Re: Revisiting recent challenges to the ancient fish-specific genome duplication hypothesis. Current Biology 11: R10071-R1008

Robinson-Rechavi, M. and V. Laudet. 2001c. Evolutionary rates of duplicate genes in fish and mammals. Molecular Biology and Evolution 18: 681–683

Robinson-Rechavi, M., O. Marchand, H. Escriva, P.L. Bardet, D. Zelus, S. Hughes, and V. Laudet. 2001d. Euteleost fish genomes are characterized by expansion of gene families. Genome Research 11: 781–788

Ronquist, F. and JP Huelsenbeck. 2003. MrBayes 3: Bayesian phylogenetic inference under mixed models. Bioinformatics 19: 1572–1574

Schluter, D. 1996. Ecological causes of adaptive radiation. American Naturalist 148: S40–S64

Schwarz, G. 1978. Estimating the Dimension of a Model. The Annals of Statistics 6: 461–464

Seehausen, O. 2006. African cichlid fish: a model system in adaptive radiation research. Proceedings of the Royal Society B-Biological Sciences 273: 1987–1998

Semon, M. and KH Wolfe. 2007. Rearrangement rate following the whole-genome duplication in teleosts. Molecular Biology and Evolution 24: 860–867

Stearley, R.F. and G.R. Smith. 1993. Phylogeny of the pacific trout and salmon (*Oncorhynchus*) and genera of the family Salmonidae. Transactions of the American Fisheries Society 122: 1–33

Stellwag, E.J. 2004. Are genome evolution, organism complexity and species diversity linked? Integrative and Comparative Biology 44: 358–365

Swofford, D. L. 2003. PAUP*. Phylogenetic Analysis Using Parsimony (*and Other Methods). Version 4. Sinauer Associates, Sunderland, Massachusetts.

Tamura, K. and M. Nei. 1993. Estimation of the number of nucleotide substitutions in the control region of mitochondrial-DNA in humans and chimpanzees. Molecular Biology and Evolution 10: 512–526

Taylor J.S., Van de Peer Y., Braasch I. and A. Meyer. 2001a. Comparative genomics provides evidence for an ancient genome duplication event in fish. Philosophical Transactions of the Royal Society of London Series B 356: 1661–1679

Taylor, J.S., Y. Van de Peer and A. Meyer. 2001b. Genome duplication, divergent resolution and speciation. Trends in Genetics 17: 299–301

Taylor, J.S., Y. Van de Peer and A. Meyer. 2001c. Revisiting recent challenges to the ancient fish-specific genome duplication hypothesis. Current Biology 11: R1005–R1007

Taylor J.S., Braasch I., Frickey T., Meyer A. and Y. Van de Peer . 2003. Genome duplication, a trait shared by 22,000 species of ray-finned fish Genome Research 13: 382–390

Taylor J.S. and J. Ráes. 2005. Small-scale gene duplications. In: The Evolution of the Genome, edited by T.R. Gregory. Elsevier, San Diego, pp. 289–327

Thompson, J.D., T.J. Gibson, F. Plewniak, F. Jeanmougin, and D.G. Higgins. 1997. The CLUSTAL_X windows interface: flexible strategies for multiple sequence alignment aided by quality analysis tools. Nucleic Acids Research 25: 4876–4882

Van de Peer I. and A. Meyer (2005). Large scale gene and ancient genome duplications. In: The Evolution of the Genome, edited by T.R. Gregory. Elsevier, San Diego, pp 329–368

Venkatesh B. 2003. Evolution and diversity of fish genomes. Current Opinion in Genetics and Development 13: 588–592

Walsh, H.E., M.G. Kidd, T. Moum, and V.L. Friesen. 1999. Polytomies and the power of phylogenetic inference. Evolution 53: 932–937

Woolfe A., Goodson M., Goode D.K., Snell P., McEwen G.K., Vavouri T., Smith S.F., North P., Callaway H., Kelly K., Walter K., Abnizova I., Gilks W., Edwards Y.J.K., Cooke J.E. and G. Elgar. 2005. Highly conserved non-coding sequences are associated with vertebrate development. PLoS Biology 3: 0116–0130

Woram, R.A., K. Gharbi, T. Sakamoto, B. Hoyheim, L.E. Holm, K. Naish, C. McGowan, M.M. Ferguson, R.B. Phillips, J. Stein et al. 2003. Comparative genome analysis of the primary sex- determining locus in salmonid fishes. Genome Research 13: 272–280

Yang Z. 1996. Maximum-likelihood models for combined analyses of multiple sequence data. Journal of Molecular Evolution 42: 587–596

Zhang, J.Z. 2003. Evolution by gene duplication: an update. Trends in Ecology and Evolution 18: 292–298

Zhu J., Sanborn J.Z., Diekhans M., Lowe C.B., Pringle T.H. and D. Haussler. 2007. Comparative genomics search for losses of long-established genes on the human lineage. PLoS Comput Biol 3: e247

